# Topographic alignment of auditory inputs to the visual cortex

**DOI:** 10.1101/2025.07.29.667237

**Authors:** Alexander Egea-Weiss, Benita Turner-Bridger, Aiste Viduolyte, Elsa Marianelli, Petr Znamenskiy, M. Florencia Iacaruso

## Abstract

Sensory cortical areas send long-range projections to cortical areas from other sensory modalities, supporting multisensory integration to generate a unified representation of the external world. However, the organizational principles underlying these extensive cross-modal connections remain poorly understood. In this study, we investigated the anatomical and functional organisation of auditory cortex inputs in the visual cortex. We found that populations of anatomically segregated auditory cortex neurons project to different visual cortical areas, broadcasting distinct auditory information to the dorsal and ventral visual processing streams. While sound frequency information was homogenously distributed across visual cortical areas, sound location information was differentially broadcast across the visual cortex. Specifically, sound azimuth and elevation were differentially encoded across visual cortical areas and streams matching the retinotopic bias of the target area. These findings suggest that cross-modal cortico-cortical connections follow a simple rule whereby specialised projection pathways are topographically aligned with the organisational principles of the target sensory area, ensuring spatially coherent integration of multisensory signals.

---

The brain integrates information from multiple sensory modalities to form a coherent representation of the external world. This integration is supported by projections between cortical networks primarily subserving different sensory modalities^1^. The organizational principles of these cross-modal projections remain largely unknown. Sensory cortical areas are hierarchically organised into primary and higher-order areas^2–4^. These higher-order areas receive inputs from populations of neurons in primary sensory areas, which exhibit specialised connectivity and functional properties^5–8^. In the visual system, higher visual areas (HVAs) are further divided into two parallel processing pathways: the dorsal and ventral streams^9,10^. In the mouse, visual cortical areas AL (anterolateral), RL(rostrolateral), A (anterior), AM (anteromedial), PM (posteromedial), have been proposed to constitute the dorsal visual stream, while areas P (posterior), POR (postrhinal), LI (laterointermediate), LM (lateromedial) constitute the ventral stream^2,10–13^. These streams are biased in their representation of the visual field, particularly with respect to elevation: areas of the dorsal and ventral streams are biased toward the lower and upper visual fields, respectively. Areas within the same stream exhibit correlated activity during development^14,15^, share feature selectivity^16,17^, and have similar input-output patterns^18–20^. The two streams have been proposed to play distinct roles in vision, such as guiding perception versus action^21,22^ or exploration versus exploitation^23^.

Both primary and higher order areas of the visual cortex (VC) receive direct projections from the auditory cortex (AC)^1,24–27^. While spatial congruency is a hallmark of multisensory integration^28^, the sound location encoded by AC axons does not align with the retinotopic representation of their local target population in the primary visual cortex (V1)^29^. Therefore, the logic of information transfer between the auditory cortex and the visual cortex remains to be deciphered. One possibility is that auditory information is uniformly broadcast across all visual cortical areas (Fig. 1a, left) providing a complete representation of auditory features across both processing streams. Alternatively, auditory projection neurons may selectively target specific sets of visual areas (Fig. 1a, right), providing functional specificity to multisensory interactions.

**Fig. 1.**
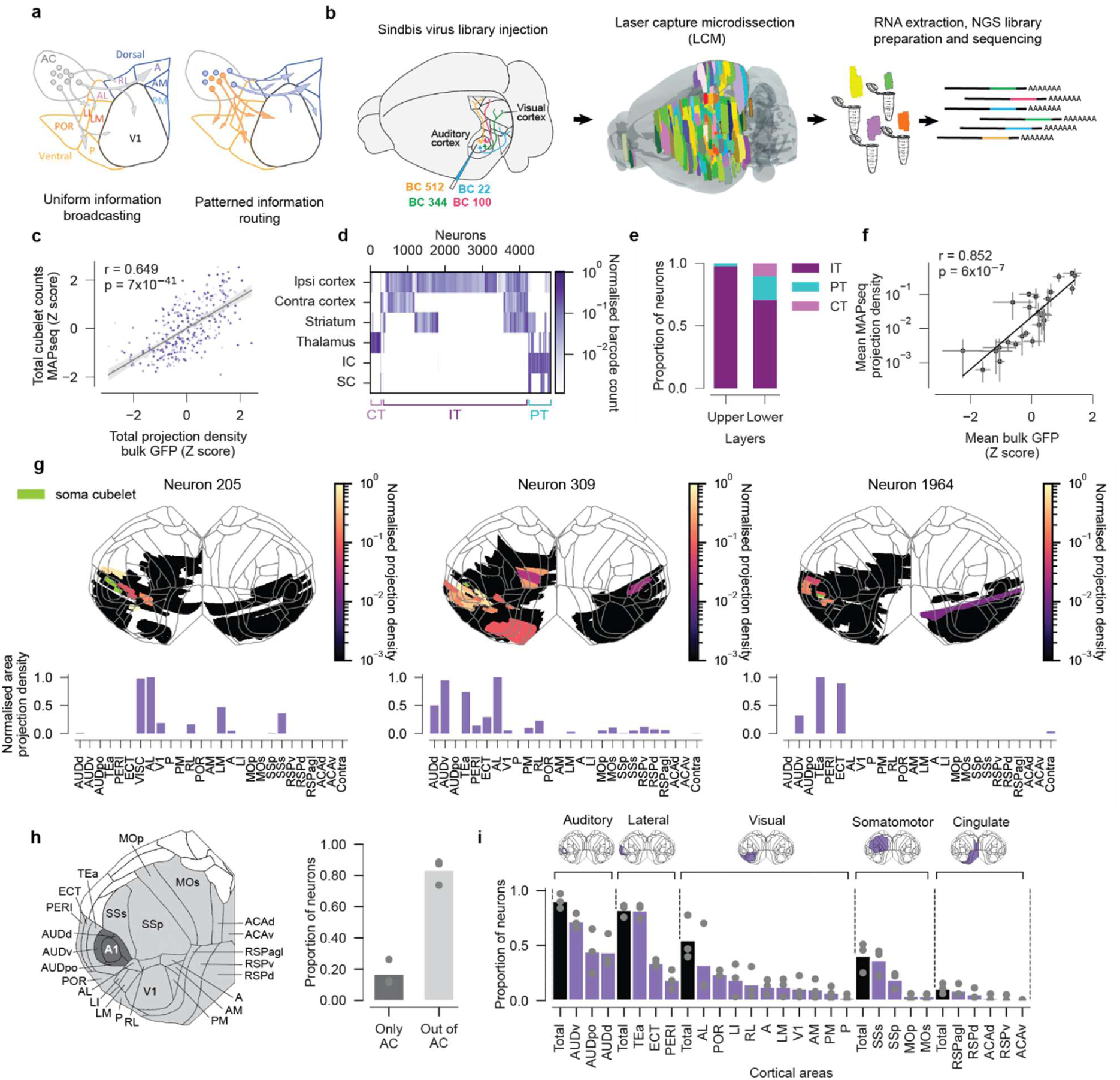
High-throughput mapping of A1 single neuron projections. **a,** Two hypothetical modes of information transfer from auditory cortex (AC) to the visual cortex. Auditory information (arrows) could be uniformly distributed across visual cortical areas (left) or selectively broadcast to specific visual areas (right). **b,** Experimental schematic for mapping single neuron projections using MAPseq. The barcoded Sindbis virus library is injected into the primary auditory cortex (A1), labelling neurons with unique RNA barcodes (BC) that are transported into axon terminals. Brain regions of interest are dissected into cubelets via laser capture microdissection (LCM) and barcodes present in each cubelet are sequenced. **c,** Total bulk MAPseq counts in individual cortical target cubelets recapitulate conventional anterograde tracing projection patterns (Pearson correlation coefficient *r* = 0.65, *p*=7×10^−42^; 338 cubelets from 3 mice, mean z-scored bulk GFP signal from 3 mice). **d,** MAPseq recapitulates brain-wide projection patterns of pyramidal tract (PT), corticothalamic (CT) and intratelencephalic (IT) neurons (4141 neurons across 3 mice). **e,** Layer distribution of IT, CT and PT neurons. CT and PT neurons are predominantly found in deep layer cubelets (885 and 1910 superficial and deep layer neurons). **f,** Mean area-level corticocortical normalised projection density estimated by MAPseq recapitulates conventional anterograde tracing (25 areas, N=3 mice for A1 MAPseq and N=3 mice for bulk anterograde tracing from A1; error bars are standard deviation across mice in log space back-transformed to linear scale; Pearson correlation coefficient r=0.85, *p*=7×10^−8^)**. g,** Example projection patterns of single neurons across cortical cubelets visualized on a cortical flatmap (top) and their projection densities across cortical areas (bottom) normalized to the maximum for each neuron. Colormap indicates normalised projection density for the sampled areas. Areas in white were not sampled. **h,** Proportions of neurons projecting only to majority auditory cortex cubelets and outside of the auditory cortex (N=3 mice). **i,** Proportions of neurons targeting different cortical areas. Black bars group neurons projecting to any of the cortical areas specified on top of the panel. Purple bars show proportions for individual areas (N=3 mice).

To distinguish between these possibilities, we analyzed the anatomical and functional organisation of individual AC neurons projecting to the visual cortex. Using high-throughput sequencing of genetically barcoded neurons (MAPseq)^30^, we found that the organization of long-range axons of auditory cortex neurons projecting to the visual cortex reflects the organization of dorsal and ventral processing streams. Specifically, AC neurons projecting to the same stream have similar co-projection patterns, and often co-project to areas within the same stream. Furthermore, we found that neurons projecting to the two streams are spatially segregated within auditory cortex. Using two-photon calcium imaging of axonal boutons from AC neurons in visual cortex, we found that these specialised projection pathways convey distinct auditory spatial information across the visual cortex, with sound location tuning of auditory inputs to the HVAs aligned to the retinotopic location of the target visual areas. These results reveal an unexpected precision of cross-modal projection pathways and suggest a simple rule underlying cortical-cortical communication, whereby cross-modal information is differentially broadcast according to the organising principles of the target area.

## Projection patterns of single primary auditory cortical neurons to the visual cortex

To determine how information flow is patterned from the primary auditory cortex (A1) to the visual cortex and surrounding cortical areas, we analysed the long-range axonal projections of thousands of neurons in parallel using multiplexed analysis of projections by high throughput sequencing^30^ (MAPseq). MAPseq relies on labelling of neurons with random RNA barcodes that are transported to their axon terminals. Projection patterns of barcoded neurons can then be reconstructed by sequencing barcodes in dissected brain regions.

To systematically characterize the projections of A1 neurons across visual cortex (VC) areas, we injected barcoded Sindbis virus into A1 and used laser capture microdissection (LCM) to dissect the ipsilateral visual cortex and adjacent areas into ∼400 x 600 µm “cubelets”. To distinguish between projections originating from superficial or deep layer neurons, we dissected the cubelets located within A1 into upper and lower layers. We also collected LCM samples from other major projection targets of the auditory cortex, including the medial geniculate nucleus in the thalamus, inferior and superior colliculus, contralateral AC, contralateral VC, and striatum. We then registered the dissected cubelets to the Allen Common Coordinate Framework^31^ and assigned neuron barcodes to different brain areas present in each cubelet^32^ (Fig.1b).

We analysed long-range projection patterns of 4,829 A1 neurons from three mice. To validate this approach, we first compared bulk MAPseq barcode counts to the projection patterns derived from fluorescent anterograde tracing using data from the Allen Connectivity Atlas^33^ (Suppl. Fig. 2a-b). Total bulk MAPseq counts were highly correlated with mean total projection density from three A1 fluorescent anterograde tracing experiments at a cubelet level across the cortex (Pearson *r* = 0.65, *p* <10^−42^, Fig. 1c), similar to the correlations observed between individual anterograde tracing experiments (mean ± S.D. = 0.75 ± 0.08; Suppl. Fig. 2c).

We classified neurons into intratelencephalic (IT), pyramidal tract (PT), and corticothalamic (CT) cell types based on their projections to the thalamus, superior and inferior colliculus. Neurons projecting to superior and inferior colliculus were labelled as PT, while the remaining neurons were classified as CT if they projected to the thalamus or IT if they did not. Corticostriatal projections in the auditory cortex originate from both IT and PT neurons^34^. While IT-type corticostriatal neurons often send axon collaterals to the contralateral auditory cortex, PT-type neurons innervate the superior and inferior colliculus and the thalamus. Our data recapitulated this segregation. Of 1,624 corticostriatal neurons, 689, 160, and 285 sent projections to the contralateral cortex, thalamus and inferior or superior colliculus, but only 3 projected both to the contralateral cortex and thalamus, and 2 projected to both contralateral cortex and inferior or superior colliculus (Fig. 1d). Thus, reconstructed projection patterns of individual A1 neurons also followed expected cell-type specific cortical wiring rules. PT, CT, and IT neurons followed the expected layer distribution patterns, with PT and CT neurons found predominantly in deep cortical layers and rarely observed in neurons originating from our superficial layer A1 cubelets (21/885 upper layer neurons, Fig. 1e). We next used single neuron projection patterns across LCM cubelets to estimate their projection density across cortical areas. Since individual cubelets were small (mean 0.22 mm^3^), we assumed uniform barcode distribution within each cubelet and distributed the projection strength between the areas contained within each cubelet according to their relative volumes (Suppl. Fig. 2d-f, see Methods). The estimated projection patterns at the area level matched those expected from bulk anterograde tracing data (Fig. 1f, Pearson’s *r =*0.85, *p*=7×10^−8^). Therefore, these data recapitulate the expected A1 connectivity patterns both at bulk and single neuron levels, confirming the specificity of MAPseq projection tracing.

Within the cortex, A1 neurons showed diverse long-range projection patterns. While some targeted cortical areas in a spatially focal manner, others exhibited more distributed projections (Fig. 1g). The majority of A1 neurons targeted the auditory cortex and nearby areas such as the temporal association area (TEa) and most (>80%) also sent projections beyond auditory areas (Fig. 1h-i). Notably, A1 neurons targeting regions within the visual cortex represented ∼50% of corticocortical projecting neurons (Fig. 1i).

## A1 projections to dorsal and ventral visual streams form distinct subnetworks

We next focused on the projection patterns of single A1 neurons in the visual cortex. VC-projecting A1 neurons most frequently targeted the POR, AL, and LI (Fig. 1i), which are closest to the A1 injection sites (Fig. 2a). Indeed, exponential decay of connection probability with distance is thought to serve as a general cortical wiring principle^35,36^. Consistent with this, we found that the frequency and normalised density of projections to cortical areas including visual areas were well fit by exponential decay with distance from A1 (p=4.3×10^−6^ and 9×10^−5^, respectively, Fig. 2b–c).

**Figure 2.**
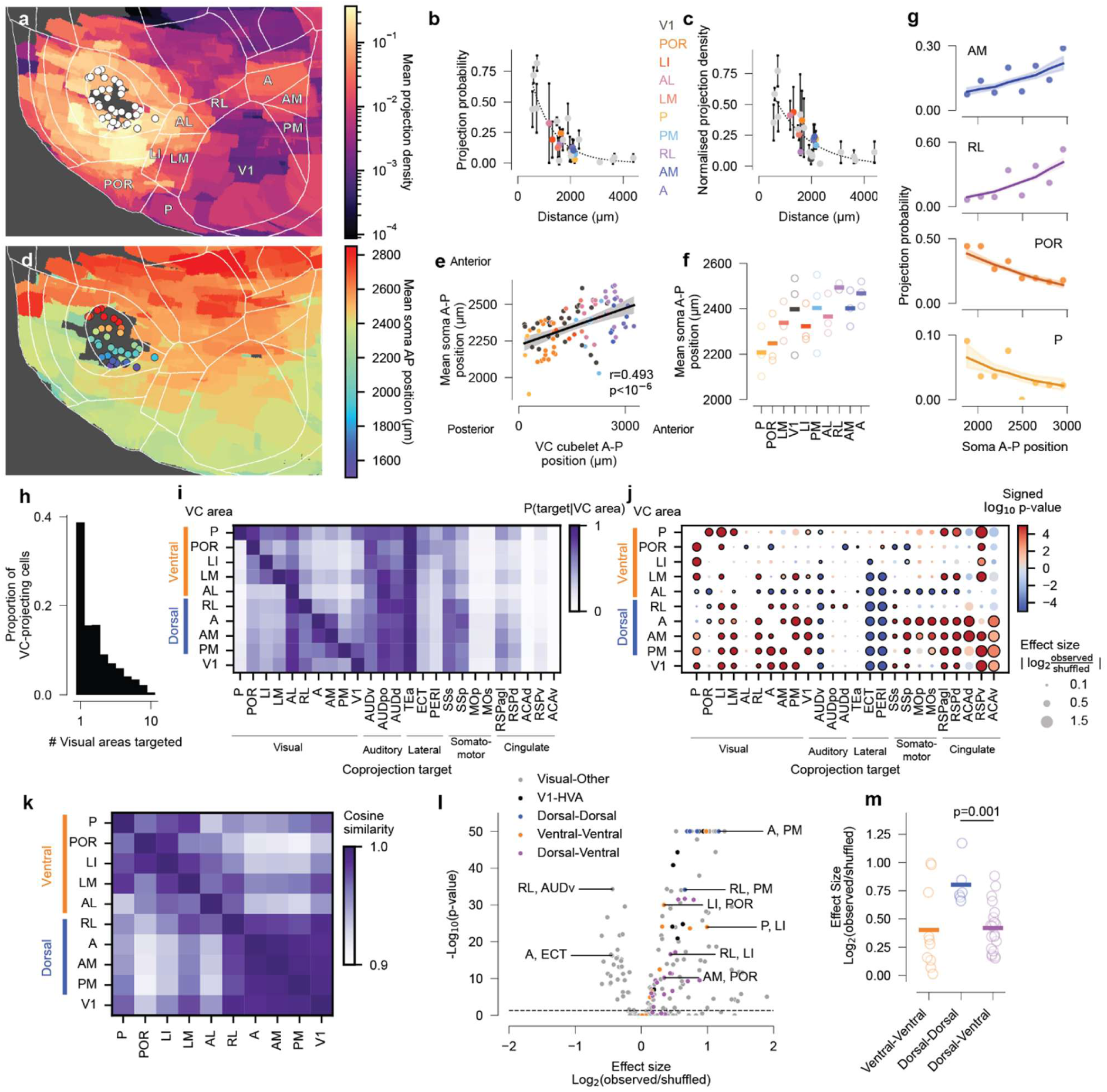
Topography of single A1 neuron projections across visual cortical areas. **a,** Cortical flatmap showing mean normalised neuron projection densities, cropped to visualise VC areas (N=3 mice). Dots indicate source site centroids. **b-c,** Targeting frequency (**c**), and normalised projection density for neurons projecting to ipsilateral cortical areas decay exponentially with distance. Dotted line shows exponential fit. **d,** Cortical flatmap showing mean A1 soma position for neuron projections to cubelets containing VC areas. Dots indicate centroid positions for A1 source site cubelets coloured according to A-P position. To minimize bias from local A1 projections and from voxels with too few neuronal projections, we analysed only cubelets containing <10% A1 and voxels containing ≥ 3 projections. **e,** A-P position for cubelets in the visual cortex plotted against mean A-P source site position for neurons targeting that VC cubelet. All values are expressed relative to the most posterior VC position. **f,** Mean normalised soma A-P position for A1 neurons targeting different VC areas. Individual points are mean for each mouse. VC areas are ordered in posterior to anterior positions **g,** Relationship between A-P source site position and targeting frequency. Y-axis is fraction of neurons targeting each plotted visual area for neurons in a particular A-P source site position. X-axis is binned A1 source site A-P position with scale in microns, values are relative to most posterior VC position. Logistic regression analysis controlling for inter-mouse variability and area-to-soma cubelet distance and adjusting for multiple comparisons (Bonferroni) showed significant effects for AM (p=6.7×10^−4^), RL (p=2.6×10^−26^), POR (p=5.6×10^−41^), and P (p=3.1×10^−8^). For visualisation, fitted lines are logistic regression model predictions for all mice together for each A-P bin position bin using the average Euclidean distance of neurons in that bin as a covariate. Shaded bands are 95% confidence interval. **h,** Histogram showing number of different VC targets of individual VC-projecting A1 neurons (N=2,647 neurons). **i,** Heatmap of conditional probability for A1 neurons targeting different cortical regions given they already project to a specific target VC area. **j**, Bubble plot corresponding to conditional probabilities shown in **i**, with circle colours indicating increased (red) or decreased (blue) conditional probabilities compared to mean shuffled population. Colorbar indicates strength of *p*-value, bubble size indicates effect size and black outline indicates *p*≤0.05 for two-tailed *p-*values compared to the null population of 100,000 shuffles and after Bonferroni correction. p-values are capped at 10^−5^. **k,** Mean cosine similarities across 3 mice for conditional probabilities across the ipsilateral cortex for A1 neurons targeting different VC areas. **l,** Volcano plot showing statistical significance of co-projection motifs involving VC areas and related log fold change over mean shuffled effect size. Here, effect size represents the number of observed co-projections between two targets in our data divided by the expected number of observing co-projection by chance. The significance values on the y-axis are capped at 50. **m,** Relative effect size compared to shuffled null population for representation of co-projection motifs (as in panel l) grouped according to streams (*p*=0.0015, Mann-Whitney test).

Because A1 is tonotopically organised along the anterior-posterior (A-P) axis^37–39^, we asked whether the projection frequency to specific VC areas depends on the A-P soma location of A1 neurons. For individual VC cubelets, their A-P position was positively correlated with the mean A-P position of somata of neurons innervating them (Fig. 2d-e; Pearson correlation coefficient = 0.49, p<1×10^−6^). On average, anterior VC areas received input from neurons located more anterior in A1 (Fig. 2f). At the single neuron level, the probability of targeting anterior visual areas A and RL was positively associated with A-P position of the neurons’ somata (Fig. 2g). Conversely, posterior visual areas POR and P were preferentially targeted by neurons in posterior A1 and the probability of targeting them was inversely related to soma position (Fig. 2g and Suppl. Fig. 4a). These data indicate that projections from A1 to the VC are topographically organised along the A-P axis. Importantly, this organisation cannot be explained by physical proximity alone since we observed a significant relationship between A-P soma cubelet position and VC areas AM, RL, P and POR after controlling for soma cubelet-to-VC area distance (Fig. 2g; p=6.7×10^−4^, 2.6×10^−26^, 5.6×10^−41^, and 3.1×10^−8^ for AM, RL, POR and P, respectively; logistic regression with mouse and distance as covariates). This trend was further confirmed from bulk anterograde tracing experiments obtained from the Allen Connectivity Atlas. We observed that axonal EGFP expression from anterior A1 injections was higher in anterior visual areas such as RL and A, while posterior A1 neurons targeted ventral areas with greater strength (Suppl. Fig. 4b-d). Together, these findings indicate that relative position along the A-P axis rather than sheer proximity alone best explains the observed topographic wiring of A1 projections across VC. The anterior VC areas AL, RL, A, AM, PM, have been proposed to constitute the dorsal visual stream, while posterior VC areas P, POR, LI, LM constitute the ventral stream^2,10^. This suggests that neurons located in anterior A1 make specialised projections to dorsal visual areas, while those in posterior A1 target the ventral stream.

We next asked whether individual A1 neurons establish dedicated projections to individual VC areas or target multiple VC areas simultaneously. We found that fewer than half (38%, 1,018/2,647) of VC-projecting A1 neurons innervated a single VC area, while the majority projected to multiple targets (Fig. 2h). This shows that similar to V1 projections^40^ individual A1 neurons broadcast information across multiple VC areas. To quantify the co-projection patterns of VC-projecting A1 neurons, we calculated the conditional probability *P*(*target*|*VC area*) of A1 neurons targeting different cortical regions given projections to a particular VC area (Fig. 2i). We compared these conditional probabilities to shuffled connectivity matrices that preserved the statistical structure of the dataset at the single area and single neuron level, including marginal projection probabilities, intrinsic biases in assignment of projections emerging from cubelet structure, and variability in barcode labelling efficiencies (Fig. 2j, Suppl. Fig. 3, see Methods).

We observed A1 neurons targeting dorsal stream areas AM, A, PM, and RL had a higher probability of co-projecting to other dorsal stream VC areas compared to the shuffled population (Fig. 2j). In addition, A1 neurons projecting to dorsal stream areas AM, A, PM were more likely to co-project to somatosensory, motor and anterior cingulate cortical areas (Fig. 2i). These projection biases were not observed for neurons innervating the ventral visual stream areas LM, LI, POR and P. Neurons projecting to both dorsal and ventral stream visual areas were less likely to co-project to the contralateral cortex and ventral auditory area (AUDv) compared to the shuffle control. Finally, neurons projecting to area AL were less likely to co-project to other visual, retrosplenial and motor cortical areas.

To quantify the similarity in projection patterns of A1 neurons targeting different VC areas, we computed the cosine similarity between their conditional probability vectors (Fig. 2k). A1 neurons targeting ventral stream VC areas exhibited co-projection patterns that were more similar to those of A1 neurons targeting other ventral stream VC areas than to those projecting to dorsal stream VC regions. The converse pattern was observed for A1 neurons projecting to dorsal stream VC areas. Interestingly, we observed projection patterns for A1 neurons that targeted AL had higher similarity to A1 co-projections involving ventral stream VC areas. This finding is consistent with recent retrograde tracing studies^41^, which demonstrated that brain-wide input connectivity of AL neurons more closely resembles that of ventral stream areas, even though AL retains features of both streams. Because of this transitional status and higher similarity with ventral stream regions in the context of single A1 neuron co-projection patterns in our dataset, we classed AL as belonging to the ventral stream in our analyses. These data demonstrate that A1 neurons projecting to the dorsal and ventral streams are organised into distinct subnetworks with heightened similarity in co-projection patterns within streams.

To further probe the organisational logic of co-projection patterns of A1→VC connections, we compared the frequency of co-innervation between pairs of cortical areas to that expected by chance given the null hypothesis that neurons project to each target region independently. This analysis identified both under- and over-represented A1 bifurcation motifs (Fig. 2k). For example, co-projections to VC areas and the ectorhinal cortex (ECT) and the ventral auditory area (AUDv) were under-represented compared to the shuffle control. On the other hand, co-projections targeting multiple VC areas were over-represented. Moreover, we found that within stream co-projections between pairs of dorsal stream areas tended to have greater over-representation compared to across stream or ventral-ventral co-projection motifs (Fig. 2l,m, p=0.0015; Mann-Whitney U test).

Together these analyses suggests that projections from A1 to the visual cortex are organised into specialized pathways, often co-projecting to multiple targets and broadcasting auditory information to specific combinations of visual areas. Neurons in posterior A1 preferentially target posterior HVAs belonging to the ventral visual stream. On the other hand, neurons in anterior A1 preferentially project to anterior dorsal stream areas and send collaterals to somatosensory, motor and anterior cingulate cortices.

## Sound frequency is transmitted uniformly to visual cortex

The anatomical segregation of A1→VC projections suggests that auditory signals might be transmitted differently to the two visual processing streams. As auditory cortex is tonotopically organised and neurons projecting to ventral and dorsal stream visual areas show different biases in their spatial location within A1 (Fig. 2d,e), we hypothesised that A1 neurons projecting to the two streams differ in their tuning to sound frequency (Fig. 3a). To test this hypothesis, we expressed the calcium indicator jGCaMP7b in auditory cortex neurons (Fig. 3b and Suppl. Fig.5a-c) and imaged their axonal projections in layer 1 of V1 and HVAs using two-photon microscopy.

**Figure 3.**
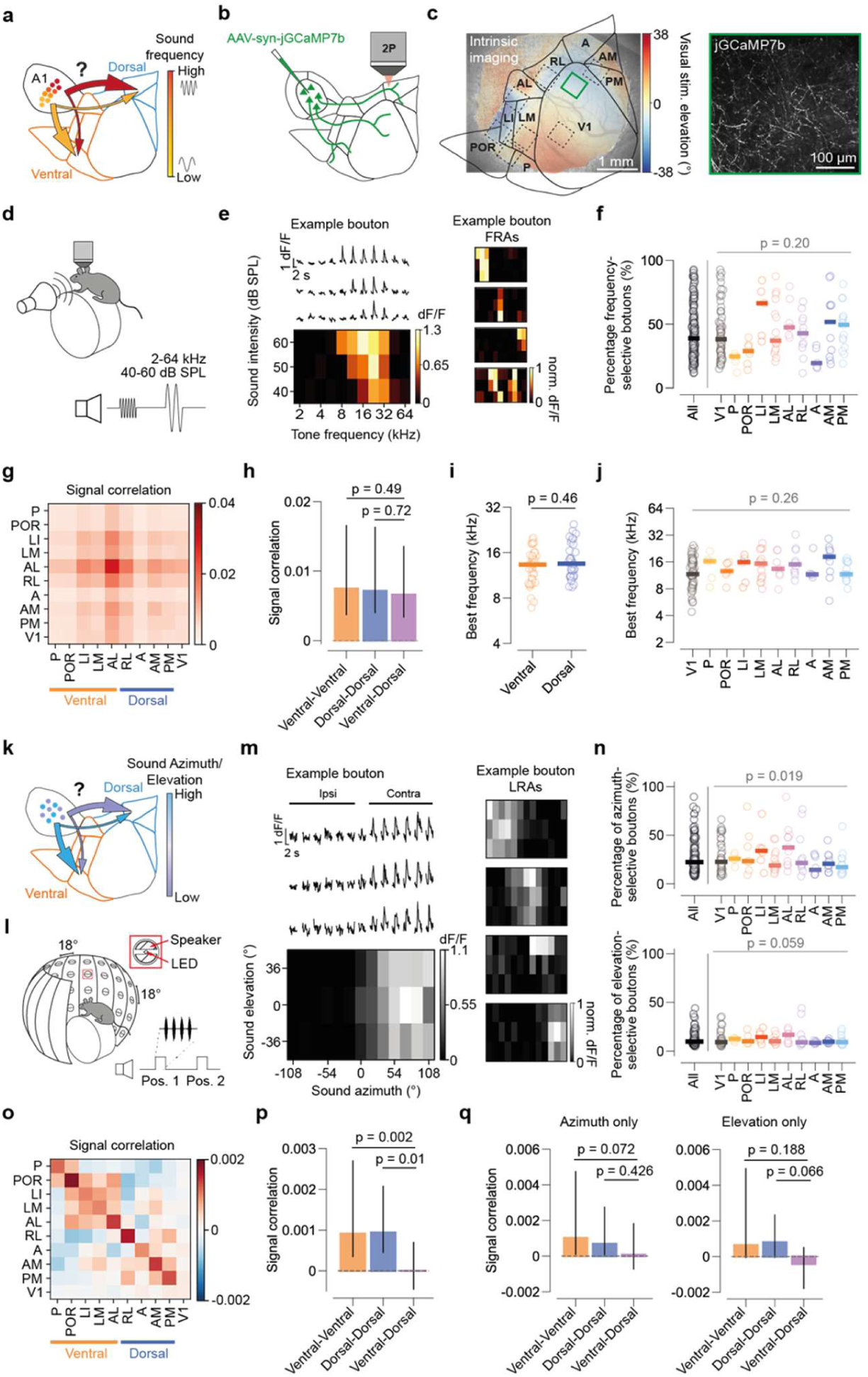
Encoding of sound location, but not frequency, differs across AC projections to visual streams. **a,** Schematic of potential tonotopic organization of auditory projections to visual streams. **b,** Schematic of experimental strategy. The calcium indicator jGCaMP7b was expressed in auditory cortex, and axonal projections were imaged in the visual cortex. **c,** Example of chronic imaging window implanted above visual cortex (left), overlaid with retinotopic map obtained with intrinsic imaging and visual area boundaries (black lines). Green square indicates location of example field-of-view (right), dashed squares indicate other recording sites from the same mouse. **d,** Schematic of auditory stimulation strategy for frequency-tuning characterisation. Head-fixed mice were presented with pure tones of different frequencies and intensities (Methods). **e,** Example fluorescence of one example AC bouton aligned to the onset of pure tone stimuli (left, top) and frequency response areas (FRAs) of same bouton (left bottom) and other example boutons (right). **f,** Percentage of tone-responsive boutons in each recording that were frequency-selective (ANOVA, p < 0.05), grouped by visual area. 1 circle = 1 recording, horizontal bar = median across recordings. Gray p-value from mixed effects ANOVA with visual area as fixed effect (Methods). N_total_ = 442648 boutons, N_responsive_ = 96539 boutons, N_freq.selective_ = 33522 boutons, 15 animals. **g,** Matrix of average pairwise signal correlations in pure-tone responses between AC boutons grouped by visual area. **h,** Signal correlation in pure-tone responses between boutons of the same or different streams. Black lines indicate hierarchically bootstrapped 95% confidence interval. Significance was assessed from the hierarchically bootstrapped difference in signal correlations (Methods). **i,** Median best frequency of boutons in each recording, grouped by stream. P-value obtained via mixed effects ANOVA with injection location and animal as fixed and random effects respectively. **j,** As in i, with recordings grouped by visual area. **k,** Schematic of potential difference in encoding of sound location (azimuth or elevation) in auditory projections to visual streams. **l,** Schematic of circular array of concentric speakers and LEDs. Speakers/LEDs were located 18° apart, spanning −108° to 108° in azimuth and −36° to 36° in elevation. Inset (bottom right) shows stimulation protocol and schematic of bandpass filtered noise bursts (Methods). **m,** Example location response areas (LRAs) of sound-location-selective AC→VC boutons. **n.** Percentages of noise-responsive boutons that were selective for sound azimuth (top) or elevation (bottom), grouped by visual area. N_total_=624087 boutons, N_responsive_ = 81559 boutons, N_azimuth.selective_ = 15556 boutons, N_elevation.selective_ = 7793 boutons, 13 animals. **o,p.** Same as **g,h**, comparing AC bouton LRAs across areas and streams. **q,** Same as **h**, for correlations in azimuth (left) or elevation (right) tuning curves between boutons of the same or different streams.

jGCaMP7b expression location within auditory cortex (AC) varied across animals and was not always confined to A1 (Suppl. Fig. 5a-c). Therefore, we hereafter refer to the population we recorded from as “AC inputs” and accounted for injection location in the analysis of functional responses by including each animal’s injection location as a factor in mixed-effects models. We first identified the location of V1 and HVAs using intrinsic signal imaging (Fig. 3b-c and Suppl. Fig. 5d). We then measured the responses of axonal boutons of AC projection neurons in these areas to pure tones of different frequencies (2-64 kHz) and sound intensities (Fig. 3d-e, 40-, 50- and 60-dB SPL). AC→VC boutons were tuned to a range of frequencies and sound intensities, and exhibited both single peaked and multi-peaked frequency response areas^42^ (FRAs, Fig. 3e). Moreover, while sound presentation increased mouse facial motion (Suppl. Fig. 6), sound frequency tuning of AC→VC boutons was largely unaffected by controlling for motor-related signals using a linear regression model (Suppl. Fig. 7). In all subsequent analyses, motor-related signals were removed from calcium traces before analysing their sound response properties. Overall, 39% of AC→VC boutons exhibited robust frequency selectivity, and the proportion of frequency-tuned boutons was consistent across all cortical visual areas (Fig 3e-f, p = 0.2, mixed effects ANOVA).

We next examined the representation of sound frequency across visual processing streams. If the representation of sound frequency by AC→VC projections segregates in a stream-like manner, auditory inputs to areas within the same stream should have more similar tone frequency tuning than inputs to areas in different streams. To test this, we assessed tuning similarity between tone-responsive boutons by computing the pairwise signal correlation of their frequency response areas (FRAs). We then averaged the signal correlations between boutons of different cortical visual areas and streams. Contrary to our predictions, the average signal correlations between areas of the same stream were not higher than those between areas of different streams (Fig. 3g-h, ventral-ventral vs dorsal-ventral, p = 0.49; dorsal-dorsal vs dorsal-ventral, p = 0.72). Indeed, the median preferred frequency did not differ between AC inputs to the two streams or across areas (Fig. 3i-j, dorsal vs ventral: p = 0.35, across areas: p = 0.26, mixed effects ANOVA). A closer comparison of the tuning properties of AC inputs to the different VC areas did not reveal differences in their representation of pure tones, including best frequency, frequency selectivity, or presence of multi-peaked FRAs, except for a broadening of the frequency tuning curves for AC→V1 compared to AC→HVAs boutons (Fig. 3j and Suppl. Fig. 8a-h). Moreover, best tone frequency did not correlate with the anatomical position of the auditory cortex injection site, suggesting that tonotopy is not reflected in AC→VC projections (Suppl. Fig. 8i-j). This is consistent with the observation that although the mouse auditory cortex is tonotopically organised at the macroscopic scale, sound frequency tuning is heterogeneous at the fine scale, showing high variability within local (∼400µm) populations^43^. Thus, the anatomical segregation of auditory inputs to the two visual streams is not reflected in their sound frequency tuning. Instead, sound frequency information is broadly distributed across visual areas.

## Sound location encoding segregates into visual streams

The auditory cortex contains spatially intermingled neurons that are tuned to sound location ^44,45^. Recent work has shown these AC-neurons provide sound-tuned inputs to visual cortex and that their inputs are not topographically organised within V1^29^. Therefore, it remains unclear whether there is any organisational logic to sound-tuned inputs to the VC. Notably, while the visual cortex is retinotopically organised, the representation of visual space is not uniform across visual areas. As a result, the dorsal and ventral streams are biased towards the lower and upper visual fields, respectively^13,46^. One intriguing possibility is that sound location information is transmitted differently to the two streams, potentially aligning with the retinotopic bias in each stream (Fig. 3j). This matching of sound and visual location could facilitate the binding of multisensory stimuli based on spatial co-localisation, as observed in subcortical regions such as the superior colliculus^47,48^. To test this hypothesis, we characterised tuning to sound location in AC→VC inputs. Mice were head-fixed at the centre of a spherical speaker array and presented with broadband noise stimuli originating from different azimuth and elevation locations, while we imaged AC→VC inputs (Fig. 3k-m). In total, 19% of sound-responsive boutons were tuned to sound azimuth, while 10% were tuned to sound elevation. The fractions of azimuth tuned boutons varied across VC areas, with boutons in areas AL and LI most frequently displaying azimuth selectivity (p_%azimuth selective_ = 0.019, p_%elevation selective_ = 0.059, mixed effects ANOVA, Fig. 3n).

To assess the similarity in sound location representation between areas, we computed pairwise signal correlations between location tuning profiles of sound-responsive boutons and grouped them by areas and streams. In contrast to the representation of sound frequency, sound location responses of boutons projecting to areas of the same stream exhibited higher signal correlations than the responses of boutons projecting to different streams (Fig. 3o-p). This trend was also present when only considering responses to different azimuths or elevations, but did not reach statistical significance (Fig. 3q, Suppl. Fig.9a). This suggests that similarities in location tuning in projections to the same stream arise through a combination of azimuth and elevation tuning and raises the intriguing possibility that azimuth and elevation preference might vary between AC axons targeting different HVAs. (Fig. 3q, Suppl. Fig.9a). Furthermore, this stream-wise organisation was also observed at the level of axons (i.e., when boutons from the same axon were grouped, see Methods, Suppl. Fig.9b), which may more closely reflect the anatomical segregation uncovered in our MAPseq experiments (Fig. 2). Thus, sound location tuning segregates in a stream-like manner, indicating that the anatomically segregated auditory projections to the two visual streams revealed with our MAPseq experiments (Fig. 2j) are functionally specialised.

## Representation of sound location in AC→VC projections

Our previous findings indicate that AC projections differentially transmit sound location information across the VC. Therefore, we separately analysed the representation of sound azimuth and elevation in AC inputs to different VC areas, as information related to these two spatial dimensions might be differentially distributed across the VC. Most AC→VC boutons were tuned to sounds on either the ipsilateral or contralateral side, with a smaller proportion tuned to central sounds, resulting in a trimodal distribution of sound azimuth tuning (Fig. 4a). Azimuth tuning curves were relatively wide, particularly in boutons tuned to ipsilateral and contralateral sounds, consistent with previous descriptions of azimuth representations in auditory cortex^44,45,49,50^ (width_ipsi-tuned_ = 105.5°; width_contra-tuned_ = 97.2°; width_centre-tuned_ = 43.7°; Suppl. Fig.9c). The shape of the azimuth tuning distribution differed between visual areas, specifically between primary and posterior visual areas (V1, P, POR) and anterior areas (RL, A, AM, Fig. 4b). Most strikingly, these areas differed in the fraction of centre-tuned boutons they receive, which was higher in dorsal stream areas than in ventral stream areas and V1 (Fig. 4d-f, dorsal vs ventral, p = 0.002, mixed effects ANOVA) Thus, inputs to V1 and posterior ventral areas were mainly tuned to ipsilateral or contralateral sounds, forming a bimodal distribution, while inputs to dorsal stream areas were more evenly distributed across ipsilateral, centre and contralateral tuning (Fig.4b-f and Suppl. Fig. 9d). In addition to differences in sound azimuth representation, AC inputs to the two streams also varied in their elevation tuning. Overall, AC→VC boutons were uniformly tuned to all elevations, with a small bias towards high elevation sounds (Fig. 4g). However, the mean preferred elevation differed between areas: AC boutons in ventral stream areas were tuned to higher elevations than boutons in dorsal stream areas (Fig. 4ih-k, dorsal vs ventral, p = 0.027, mixed effects ANOVA). Thus, AC inputs encode both sound azimuth and elevation differently depending on their target visual stream. Notably, location tuning in AC→VC boutons did not differ between animals with anterior and posterior injection sites (Suppl. Fig. 10a-g), indicating that stronger projections from anterior and posterior AC to dorsal and ventral streams are unlikely to account for differences in location tuning. Interestingly, there was a relationship between the representation of sound azimuth and dorso-ventral injection location, ventral-injected animals having fewer centre-tuned boutons than dorsal-injected animals (Suppl. Fig. 10h-n). This difference did not account for our earlier results, since dorso-ventral injection location was incorporated into all linear mixed models, and differences in percentages of centre-tuned boutons across areas were present in both dorsal-and ventral-injected animals (Suppl. Fig. 10k).

**Figure 4.**
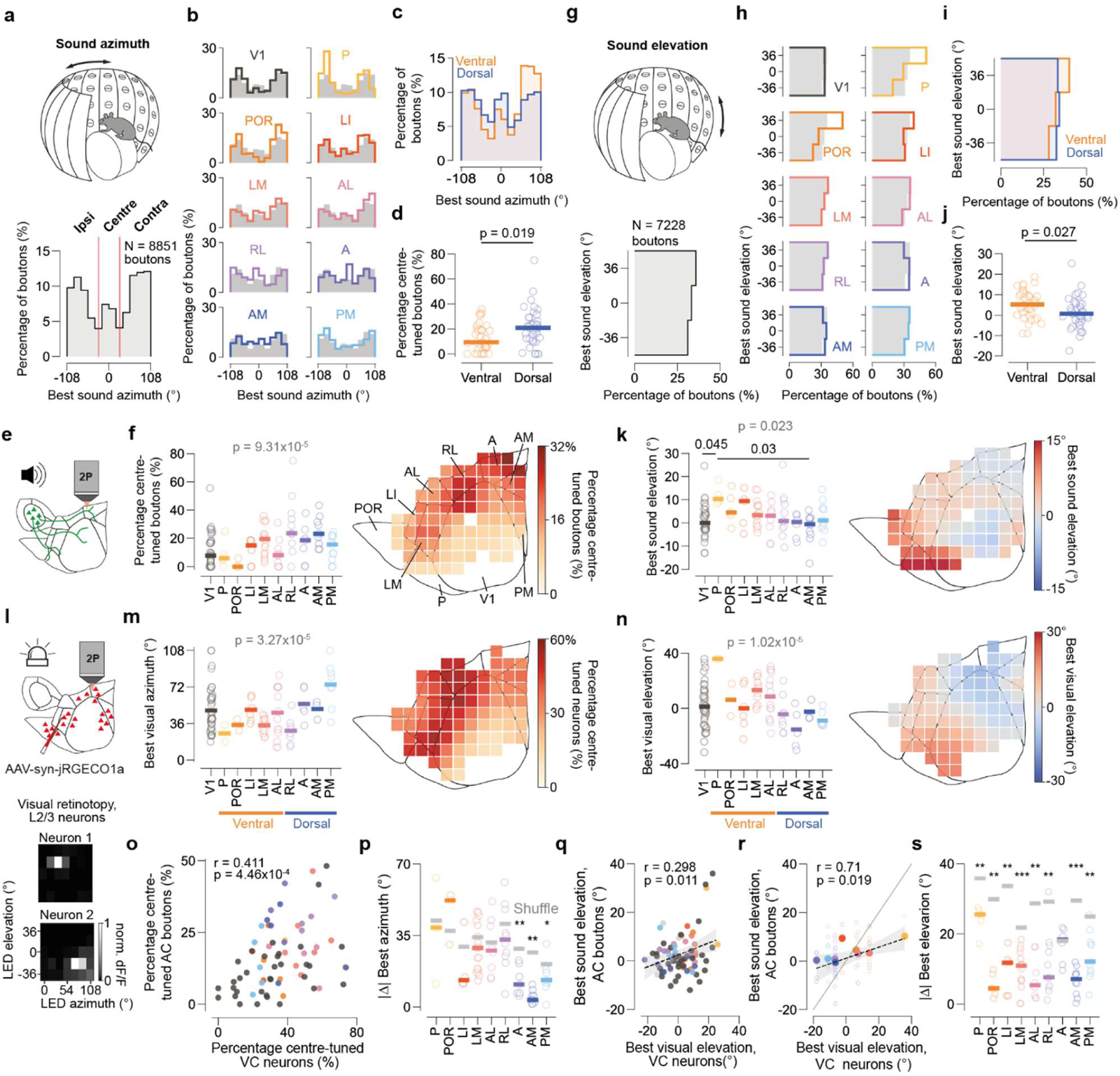
Location tuning in AC→VC inputs is biased towards retinotopy of target HVAs. **a,** Distribution of best sound azimuth for all azimuth-selective boutons that were well fit with a Gaussian (*r*^2^ > 0.6). Boutons were subsequently grouped into ipsilateral, centre or contralateral tuned (red lines). **b,** Distribution of best sound azimuth for AC boutons in different visual areas (colored). Gray distributions are for all boutons combined. **c,** Distribution of best sound azimuth for AC boutons in dorsal and ventral streams. **d,** Percentage of centre-tuned (between −30° and 30°) boutons in the two visual streams. Each circle corresponds to one session, thick lines indicate median across sessions. P-value obtained from linear mixed model. **e,** Schematic of two-photon imaging of auditory projections to visual cortex. **f,** Left, Percentage of centre-tuned boutons per recording, grouped by visual area. Each circle is one recording, horizontal bars indicate median across recordings. Gray p-value indicates comparison across all areas (mixed effects ANOVA). Right, percentage of centre-tuned boutons per spatial bin (300 um/bin), aligned across animals. Gray color (dotted line in scale-bar) indicates chance level. **g,** Distribution of best elevation for all elevation-selective boutons. **h**, Distribution of best sound azimuth for AC boutons in different visual areas (colored). **i,** Distribution of best elevation for boutons in the dorsal and ventral streams. **j,** Best sound elevation per recording for boutons in dorsal and ventral streams. P-value obtained from linear mixed model. **k,** Left, mean best sound elevation per recording, grouped by visual area. Gray p-value indicates comparison across all areas (mixed effects ANOVA), black p-values indicate comparisons between area pairs (Mann Whitney U test). Right, average best sound elevation per spatial bin. **l,** Top, schematic of viral injections to express jRGECO1a in visual cortex neurons. In some cases, both indicators were expressed in the same animal. Bottom, example responses to flashing LED stimuli in different locations of two VC neurons. Retinotopy was only mapped on the contralateral side. **m,** Left, average best visual azimuth for VC neurons per recording, grouped by visual area. Right, percentage of centre-tuned (< 30°) VC neurons per spatial bin on aligned cortical map. **n,** same as **m**, for best visual elevation in VC neurons. **o**, Correlation between percentage of centre-tuned VC neurons and percentage of centre-tuned AC boutons in each spatial bin. R and p value obtained with Spearman correlation. **p,** Absolute difference (Δ) between best visual azimuth of each area (VC neurons) and the best sound azimuth of each recording (AC boutons). Colored bars indicate median distance per area, gray bars indicate distance after shuffling AC boutons across areas. Asterisks indicate significant difference between measured distances and shuffle controls (Mann Whitney U test, *: p< 0.05, **: p < 0.01, ***: p < 0.001). **q,** Correlation between best visual elevation in VC neurons and best sound elevation in AC boutons in each spatial bin. R and p value obtained with Pearson correlation. **r**, Relationship between average best visual elevation VC neurons and best sound elevation in AC boutons across HVAs. R and p-value obtained with Pearson correlation. **s**, same as **p**, for absolute difference in stimulus elevation.

The presence of biased representations of sound location in AC→VC projections raises the question of whether these biases align with the retinotopic biases of the target areas. To evaluate this possibility, we first injected the red calcium indicator jRGECO1a into the visual cortex and mapped visual retinotopy at the single neuron level in V1 and HVAs (Fig. 4l). As expected, retinotopic maps obtained from L2/3 neurons in HVAs exhibited biases in their representations of azimuth and elevation^12,13^ (Fig. 4m,n). We then compared the representations of visual space in VC neurons and auditory space in AC boutons.

To compare the representation of stimulus azimuth in these neural populations, we focused on the relative representation of frontal (i.e. central) stimuli, as this feature varied most strongly across the AC projections to the different visual areas. The percentage of centre-tuned AC boutons correlated positively with the percentage of centre-tuned VC neurons across the visual cortex (r = 0.41, p = 4.46×10^−4^, Spearman correlation, Fig. 4o).

This positive correlation was also found within the primary visual cortex (Suppl. Fig. 9e, r = 0.49, p = 0.004), and suggests that the proportion of centre-tuned AC inputs to a given region of VC is tailored to its retinotopic preferences. Notably, we found no consistent relationship between the best sound azimuth in AC boutons and best visual azimuth in VC neurons, including within V1, consistent with previously published findings^29^ (Suppl. Fig. 9f). However, we did observe that the difference (Δ) between preferred sound azimuth in AC boutons and mean preferred visual azimuth in VC neurons in areas A, AM and PM was lower than in the respective shuffle controls (Fig. 4p), suggesting that AC inputs to medial visual areas are aligned with the retinotopic preferences of their targets. The representation of stimulus elevation in AC boutons and VC neurons was also correlated, both at the level of the whole VC and across higher visual areas (Fig. 4q,r). Accordingly, the mean distance between the preferred sound elevation in AC boutons and the preferred visual elevation in VC neurons was lower than in a shuffle control in all HVAs except for area A (Fig. 4s). Thus, auditory inputs to the VC convey auditory spatial information that is aligned with the retinotopic preferences of their target areas.

## Discussion

Our results reveal a fundamental principle of cortico-cortical communication, whereby cross-modal information is differentially broadcast according to the organising principles of the target area. We found that auditory cortex projections to visual cortex are anatomically segregated, following the division of the visual cortex into two processing streams. Neurons in anterior AC preferentially target dorsal stream areas while neurons in posterior AC target ventral stream areas. Moreover, individual AC neurons exhibit stream-specific co-projection patterns, preferentially targeting multiple areas within the same stream. This anatomical segregation at the population and single-neuron level results in differential information transfer from auditory cortex to visual cortex, such that auditory spatial information is differentially distributed to the two visual processing streams. Furthermore, the auditory spatial information in AC→VC projections is biased towards the retinotopic location of their target, thus aligning with the intrinsic functional organisation of the visual cortex.

The differential representation of specific cortical co-projection motifs was previously found in V1→HVA projections^40^, suggesting that specificity in co-projection patterns may be a common feature of cortico-cortical communication. These results indicate that similar rules may govern the co-projection patterns of V1 and AC neurons across HVAs. Bifurcating AC→VC projections provide a mechanism for broadcasting the same auditory information within, but not across, visual processing streams. Such structured connectivity at the stream level could ensure specificity while facilitating parallel processing of multisensory information across functionally distinct visual areas within each stream. AC neurons projecting within a stream also exhibit greater similarity in their co-projection patterns to other cortical targets beyond the visual cortex. This suggests that these specialised output channels are likely to extend to other sensory modalities providing the basis for unified cortical multisensory representations.

Our finding that sound-location tuning in AC inputs is spatially biased towards the retinotopy of their target visual area is consistent with the organisation of feedback projections within the visual cortex, which tend to preserve local retinotopy^51–53^. Furthermore, our findings align with the principle of spatial coincidence as a hallmark of multisensory integration, extensively documented in subcortical structures such as the superior colliculus^28,54^. The relatively coarse alignment of auditory tuning with retinotopic location may reflect the broad spatial receptive fields of auditory cortical neurons compared to the superior colliculus^44,45,49,50^. Interestingly, our results provide functional evidence for auditory biases along the elevation axis between the dorsal and ventral streams^12,13^, extending visual stream-specific elevation differences into the auditory domain. Further supporting the role of retinotopic bias in cortical multisensory interactions, neurons in area RL encode spatially coincident visuo-tactile information from proximal space^55,56^ and contribute to the generalisation of learned associations across sensory modalities^57^. Consistent with this, our data show that RL receives auditory inputs biased to centre azimuths and middle elevations. These results beg the question of how cross-modal cortico-cortical projections to higher sensory areas contribute to multisensory perception and behaviour, particularly in tasks requiring spatially aligned auditory and visual processing.

Despite the anatomical segregation of AC neurons projecting to the dorsal and ventral streams, we found no evidence for tonotopic organisation of AC→VC projections. This may be due to the widespread expression of our calcium indicator, which may cover multiple tonotopic regions in the AC, the local salt-and-pepper organisation of frequency tuning in AC^58^, and the challenges of measuring global tonotopic organisation from supra-threshold responses in the mouse AC^43^. Together, our MAPseq and functional results suggest that the representation of sound location may vary across different anatomical regions of the auditory cortex, for example between dorsal and ventral auditory cortex (Suppl. Fig.10), suggesting that the auditory cortex contains a map of auditory space. However, no physiological study to date has identified such a map. Instead, sound-source locations are thought to be represented by widely distributed populations of broadly tuned neurons^45,59^. Our findings raise the possibility that a cortical map of auditory space could emerge in the AC within the subset of sound location-encoding neurons that project to the VC.

The anatomically structured and functionally specialised cross-modal projections to the visual cortex we identified may develop under the influence of visual experience, analogous to the role of visual retinotopy in aligning visual and auditory maps in the superior colliculus during development ^47,60^. The visual critical period overlaps with the development of binaural integration^61^, and direct VC→AC projections have been proposed to gate auditory critical periods by regulating the impact of visual experience on auditory processing^62^.

Together, our results demonstrate that multisensory cortico-cortical communication follows the functional organisation of the target sensory system. While sound location information is functionally segregated, the uniform representation of sound frequency across visual areas may ensure that all visual regions receive a complete auditory spectral representation, facilitating robust integration of auditory signals into visual processing. This suggests that during multisensory behaviour, cross-modal cortico-cortical projections may support perceptual processing of visual and auditory stimuli originating from the same spatial source, providing a mechanistic basis for multisensory integration in higher cortical visual areas.

## Methods

### Animals

All animal procedures were licensed by the UK Home Office and approved by the Crick Institutional Animal Welfare Ethical Review Panel (PEB4A5081 & PP2817210). Experiments were performed on a total of 21 mice (male and female) of the C57BL/6J.Cdh23753A>G^63^ (MRC Harwell, UK) strain, aged 2-4 months, including 3 mice used for MAPseq experiments and 18 mice used for calcium imaging experiments. Mice were bred and maintained at the mouse facility of the Francis Crick Institute, with controlled temperature (21 ± 2 °C) and humidity (55 ± 10 % RH), with *ad libitum* access to food (Teklad global diet, Envigo, UK) and water, and kept on a 12 h:12 h light/dark cycle (lights on at 10 pm).

### Surgeries

24 hours before surgery, mice were given an analgesic (Metacam in custard, 1.5 mg/ml, 1:1 Metacam: custard). Mice received a subcutaneous injection of buprenorphine (0.1 mg/kg) and meloxicam (10 mg/kg) immediately before surgery. Mice were then anaesthetised with isoflurane (1%-5%), and their heads were shaved. They were then placed on a heating pad under a stereotaxic frame (Kopf Instruments). The scalp was cleaned using a chlorhexidine solution (0.8%). The mouse’s temperature was maintained at 35-37°C during surgery using a DC temperature controller (FHC) with a rectal thermometer input and a heating mat output. Anaesthesia depth was monitored by observing the respiratory pattern. The mouse’s eyes were covered with eye ointment (Maxitrol) to prevent desiccation. All injections were performed using glass microinjection needles, using a pressure microinjector (Nanoject III, Drummond). All surgeries were performed on the left cranial hemisphere.

For MAPseq experiments, the scalp was incised with a scalpel, and the temporal muscle was detached from the skull to expose the squamosal bone. The animal’s head was tilted clockwise by ∼20° along the roll axis, and the microinjector was angled at 20° counterclockwise to ensure that the injection was performed nearly perpendicular to the brain surface. The MAPseq Sindbis virus library (JK100L2, MAPseq/BARseq Core facility, Cold Spring Harbor Laboratory) was injected through a small craniotomy into the auditory cortex. Each animal received 3 injections, placed 0.8, 1.2 and 1.6 mm anterior to the juncture of the parietal, interparietal and squamosal skull bones, and dorso-ventrally positioned just below the temporal muscle’s attachment point to the skull. Injection depth ranged between −500 and −250 µm from the pia, with 150 nl per injection (100 nl at – 500 µm and 50 nl at –250 µm, injection rate: 1 nl/s). After injections, the craniotomy was sealed with VetBond (3M), and the scalp was sutured.

For *in vivo* imaging experiments, mice received an intraperitoneal injection of dexamethasone (2-3 mg/kg), 3-5 hours before surgery. A portion of the scalp was removed bilaterally, access to the auditory cortex was obtained by detaching the temporal muscle and rotating the head as described above. 100 nl of AAV1-hSyn-jGCaMP7b (Addgene) were injected 1.4-1.6 mm anterior to the juncture of the parietal, interparietal and squamosal skull bones, at a depth of 500 µm. The head was then rotated back to a horizontal position, and a custom-made circular metallic head-fixation plate was attached to the skull above the visual cortex using dental cement (SuperBond C&B, Sun Medical). A circular craniotomy (4 mm diameter) was performed above the visual cortex using lambda as a landmark; the skull was removed, and the dura was carefully resected with forceps. In some animals (11/18), to allow imaging of neural activity in the visual cortex, 5 to 8 injections of AAV1-hSyn-jRGECO1a (Addgene) were made across the visual cortex, using blood vessels as landmarks.

For these injections 50-70 nL were delivered at each location, at ∼250 µm depth. In all animals, the craniotomy was covered with a glass coverslip (4 mm diameter), and secured with VetBond and dental cement. Animals were allowed to recover for a week before intrinsic imaging. Two-photon imaging commenced 3 weeks after injections.

### Spike-in RNA

Spike-in RNA was *in vitro* transcribed from Ultramer® duplex oligo (Sequence 5’-3’: ATGATCATAATACGACTCACTATAGGGGACGAGCTGTACAAGTAAACGCGTAATGATACGGCGACCACCGAGATCTACACTCTTTCCCTACACGACGCTCTTCCGATCTNNNNNNNNNNNNNNNNNNNNNNNNATCAGTCATCGGAGCGGCCGCTACCTAATTGCCGTCGTGAGGTACGACCACCGCTAGCTGTACA, Integrated DNA Technologies) as described previously^30^ using MEGAscript T7 Transcription kit (Invitrogen, #AM1333) for MAPseq optimisation, and T7 MEGAshort kit (Invitrogen, #AM1354) for A1 MAPseq experiments. We included the DNAse treatment step after *in vitro* transcription to remove template DNA. The *in vitro* transcribed RNA was purified using the RNeasy Mini Kit (QIAGEN, #74106) and stored in aliquots at −80°C.

### MAPseq tissue preparation and laser capture microdissection

Two days after injection, mice were culled by intraperitoneal injection of pentobarbitone (1.4 mg g^−1^), the brain was dissected and flash frozen in Optimal Cutting Temperature (OCT) compound within a cryomold over 100% ethanol on dry ice and stored at −80 °C. Coronal brain sections (200 μm thick) were cryosectioned onto PEN-membrane slides (Leica Microsystems #11600289). Immediately after mounting, sections were fixed for 3 min at 4 °C with ice-cold ethanol. Sections were then rinsed with ice-cold water and stained twice in ice-cold 0.5% toluidine blue for 30 s (Thermo Scientific, #348600250). Samples were rinsed three times in ice-cold water, fixed twice in ice-cold 75% ethanol at 4 °C for 2 min, and dried in a vacuum desiccator for 30 mins at room temperature. The samples were then stored at −80 °C in a 50 ml falcon tube with desiccant. Laser capture microdissection (LCM) was performed using the Leica LMD 7000. Cortical areas of interest were dissected into approximately 600 μm arc-length cubelets. In addition, subcortical targets of the auditory cortex, including thalamus, striatum, and tectum, as well as regions of the olfactory bulb as a negative control, were dissected by LCM. Equivalent areas in adjacent sections were typically pooled generating cubelets approximately 400 μm thick. The primary auditory cortex was further dissected into upper and lower layers of approximately 300 μm thickness for each half. For all LCM samples, dissections aimed to avoid regions containing fibres of passage. Samples were collected into 0.5 ml tubes (Greiner Bio-One, #682201) containing 65 μl of buffer RLT (QIAGEN, #74004) with 40 mM Dithiothreitol (DTT, Promega, #V3151), placed on dry ice and stored at −80 °C. LCM samples were homogenized by incubating for 5 min at room temperature in buffer RLT with DTT (350 μl total volume), combined with physical disruption using a pipettor. RNA was extracted using RNeasy Micro kit (QIAGEN, #74004) without the DNA digestion step and eluted in 14 μl water. RNA quality was assessed using a TapeStation D1000 (Agilent) on 1 μl of each sample.

### MAPseq sequencing library preparation

PCR primer sequences for MAPseq library preparation were adapted for compatibility with the NovaSeq sequencing platform. For each reverse transcription reaction, 8 μl of RNA eluate per LCM sample was combined with 1×10^−7^μg/μl spike-in RNA, 1 μl 10mM dNTPs (Thermo Scientific, #10319879), 2 μl water, and 1 μl of 10 μM reverse transcription primer (MAPSEQ_NOVA_RT### in the primer table list, the oligo sequence: TGACTGGAGTTCAGACGTGTGCTCTTCCGATCTNNNNNNNNNNNNNNXXXXXXXXXXXXXXXXTGTACA GCTAGCGGTGGTCG contains a unique 16 nucleotide barcode sequence (X_16_), enabling assignment of sequencing reads to different LCM samples, as well as a random 14 nucleotide UMI). The reaction was heated to 70 °C for 10mins, placed on ice for 5 mins, and 4 μl Superscript IV buffer, 1 μl 100mM DTT, 1 μl RNaseOUT (Thermo Fisher Scientific, #10777019), 1 μl SuperScript IV reverse transcriptase (Thermo Fisher Scientific, #18090050) were added to the RNA mix and incubated at 55 °C for 50 minutes and 80 °C for 10 minutes. 1μl RNAse H (New England Biolabs, #M0297L) was added and incubated at 37 °C for 20 minutes. Second strand synthesis was then performed, adding 1.5μl of 10 μM targeted second strand synthesis primer (MAPseq_NOVA_SSS_006), 5μl 10X reaction mix, 0.4μl AccuPrime Pfx DNA polymerase (Thermo Fisher Scientific, #12344032) and 22.1μl water, the reaction was heated to 95 °C for 2 mins 15 sec, annealed at 55 °C 1 min, extended 68 °C 1 min. All samples were then pooled together, and bead clean-up performed using KAPA Pure Beads (Roche, #KK8001) using 1.8x beads ratio then eluted in 10 μl water per number of samples. 80% of the eluted pooled second strand synthesis product were then split across 96 well PCR plates. Per well, 8 μl of the double stranded cDNA eluate was exonuclease treated by adding 1 μl Thermolabile Exonuclease I (New England Biolabs, M0568L) and 1 μl of buffer r3.1 and incubated in a thermocycler for 30 mins at 37 °C followed by 5 mins at 80 °C. ExoI-treated cDNA was then amplified by nested PCR. First, 10 μl of the ExoI reaction was amplified using Accuprime pfx polymerase (Thermo Fisher Scientific, #12344032) and primers MAPseq_NOVA_SSS_006 and MAPseq_NOVA_PCR1_007 in a 50 μl volume. PCR cycle conditions were: 2 mins at 95 °C, followed by 20 cycles of 95°C for 15 seconds and 68°C for 1 minute. 10μl Thermolabile Exonuclease I (New England Biolabs, M0568L) was then added to the PCR reaction and incubated for 30 mins at 37 °C followed by 5 mins at 80 °C. Samples were pooled again and 1/3 of the PCR product was purified using KAPA Pure Beads (Roche, #KK8001) with 1.5x bead ratio and eluted in 11 μl water multiplied by number of samples and split across 96 well plates using the same number of wells as there were samples. In each well, 10 μl of the eluate was used in a second PCR reaction (primers: MAPseq_NOVA_PCR2_008, MAPseq_NOVA_PCR2_009) using AccuPrime pfx polymerase (Thermo Fisher Scientific, #12344032), PCR cycle conditions: 2 mins at 95 °C followed by 8 cycles of 95 °C for 15 seconds and 68 °C for 1 minute. The final PCR product was purified using Monarch PCR & DNA Clean-up Kit (New England Biolabs, # T1030L) and run on a 1.5% agarose gel. The ∼250bp band was gel extracted using Wizard SV Gel and PCR Clean-Up System (Promega, #A9281). DNA sequencing was performed with 500 million reads per mouse using NovaSeq 6000 or NovaSeq X (Illumina). For one mouse (FIAA55.4d), 9μl RNA was used per LCM sample (rather than 8μl) in the 1^st^ strand synthesis reaction, together with a 10% increase in second strand synthesis eluate used after bead clean-up and corresponding increase in number of PCR reactions.

### Registration of laser capture microdissection cubelets to the Allen Common Coordinate Framework (CCF)

To register LCM cubelets to the Allen Common Coordinate Framework (CCF) v3^31^, overview images of the section were acquired before LCM and after each LCM cubelet had been dissected. Each post-LCM cubelet image was aligned with the original section overview image using a custom MATLAB script through automated SURF-based feature mapping, or control-point registration if automated feature mapping was not successful. Cubelet boundaries were manually annotated on aligned pre- and post-LCM images.

The overview coronal sections were registered to the Allen CCF v3^31^ using QuickNII and VisuAlign software^64^. These data were then used to transform each pixel in each LCM slice into Allen CCF coordinates in Python. To calculate the Allen CCF coordinates for each LCM cubelet, we then used the binary mask to extract Allen CCF coordinates from each 3D coronal slice and annotate every coordinate in the LCM sample to with the corresponding coordinates in the 25μm resolution Allen CCF volume. Larger non-linear deformations – particularly those arising from separation of cortical hemispheres across the midline – led some pixels to be erroneously assigned to the contralateral hemisphere, as well as occasional missing pixel assignments. We corrected these artifacts by removing erroneous contralateral assignments and filling any holes contained within LCM sample annotation.

### MAPseq sequencing data pre-processing

All barcode sequencing data analysis was performed in Python. Raw barcode counts were comprised of two fastq files each with 100bp reads for either end of the MAPseq PCR amplicon. These reads were combined and filtered to select reads containing the expected consensus sequence GCGGC at the positions 37 to 42 of the read. Reads were assigned to the LCM samples they originated from with *fastx_barcode_splitter,* using the unique 16-nucleotide sample barcode sequence specified in the gene-specific reverse transcription primer, allowing for a maximum of two mismatches. We corrected for sequencing and PCR errors for the 30 nucleotide neuron barcode sequences in each sample using the UMI-tools API^65^ employing the directional clusterer approach. Subsequently, the 14 nucleotide UMI sequences for each neuron in each sample were similarly corrected for sequencing/PCR errors using directional clustering in the UMI-tools API. To further reduce the likelihood of including reads arising from sequencing errors, we applied a minimum UMI-duplicate count threshold for neuron barcode reads. Thresholds were determined by visually inspecting UMI-duplicate count histograms for each mouse dataset, which showed a bimodal distribution with a smaller peak at low UMI counts indicative of potential spurious reads. The cut-off was chosen to exclude this low-count noise population. Consequently, cut-off values of 5, 4, and 2 UMI duplicates were used for mice FIAA45.6a, FIAA45.6d, and FIAA55.4d, respectively.

Template switching has previously been reported as a complication arising from pooling multiple samples during the MAPseq library preparation^32^. This occurs during PCR amplification when the DNA polymerase detaches mid-elongation, and the truncated product subsequently anneals to a different template strand that shares the same sequence, resulting in chimeric PCR amplicons^66^, Suppl. Fig. 1c). To correct for template switching events, we exploited the high sequencing depth made achievable by adapting MAPseq for the high-throughput NOVAseq platform. We firstly identified instances where the same UMI sequence was shared by different neuron barcodes. We reasoned that shared UMI sequences across neurons could be due to: (i) re-use of the same UMIs from biases in random nucleotide selection during oligo synthesis, (ii) low quality reads with low complexity sequences caused by sequencing artifacts, (iii) template switching events. Since template switching events are a rare occurrence during a PCR amplification cycle^66^, the original neuron-barcode/UMI combination would be expected to be present in much higher abundance compared to the template switched neuron-barcode/UMI combination. For each re-used UMI we therefore calculated the ratio of the neuron-barcode/UMI duplicate count for the most abundant neuron-barcode/UMI combination (1^st^ max) and compared it to the next most abundant neuron-barcode/UMI duplicate count (2^nd^ max). For 1^st^ max/2^nd^ max ratios of less than 10, we observed an increase in AT-content and a decrease in entropy for UMI sequences (Suppl. Fig. 1d), suggesting these were likely due to factors (i) or (ii). Consequently, shared UMIs with a 1^st^ max/2^nd^ max ratios below 10 were discarded, while shared UMIs with ratios ≥ 10 were considered template switching events. In these cases, we retained the 1^st^ max neuron-barcode/UMI combination while discarding all the neuron-barcode/UMI combinations that were less abundant.

Next, all neuron barcode UMI counts were deduplicated and separated from barcodes corresponding to spike-in RNA by the spike-in specific barcode sequence: N_24_ATCAGTCA. Finally, neuron barcode counts per LCM sample were collated into a single matrix.

### MAPseq data filtering, normalization and soma identification

Neuron barcodes that were present at less than 100 barcode counts across the whole dataset were discarded. Comparing barcode counts in A1 target samples to the OB negative control samples showed the vast majority (95%) of barcodes in the negative control were present as single counts (Suppl. Fig. 1e). We therefore considered this background and only included barcode counts in a sample if present at ≥2 counts. Next, as a quality check to identify LCM samples with poor RNA yield, samples where the total MAPseq neuron barcode count was zero were removed. Barcode counts were then normalised in two steps. First, for each sample we calculated a scaling factor by dividing that sample’s spike-in RNA count by the median spike-in count across all samples. Next we divided MAPseq neuron barcode counts in each sample by that sample’s scaling factor, thereby correcting for variation in library preparation efficiency across samples.

To subsequently identify which samples contained the neuron’s soma for each barcode, we took a similar approach to BRICseq^32^. For every neuron barcode we identified the sample containing the maximum amount of that barcode (soma_max_). We then identified the sample containing the next largest number of counts for that barcode (2^nd^_max_), excluding samples adjacent to the soma_max_ sample in 3D space as they were likely to contain dendrites and local axonal projections. We considered soma_max_ to be a soma-containing sample only when the soma_max_/2^nd^_max_ count ratio was ≥5. Barcodes where soma_max_/2^nd^_max_ count ratio <5 were discarded. We then filtered for neurons that contained a minimum of 10 barcode counts in the 2^nd^_max_ sample as well as selecting for neurons where soma_max_ LCM cubelet contained predominantly A1.

### Comparison of A1 MAPseq to anterograde tracing datasets

To compare mesoscale anterograde tracing datasets from A1 to the A1 MAPseq dataset, we used EGFP anterograde tracing data from the Allen Mouse Brain Connectivity Atlas^33^. For comparison of cortex-wide projection strength patterns (Fig. 1c and 1f as well as Suppl. Fig. 2b-c), we selected experiments where EGFP-expressing AAV was injected into wild-type mice and ≥75% of the injection site was in A1 (experiment IDs: 120491896, 116903230, 100149109; available from connectivity.brain-map.org/). For the comparison of anterograde tracing experiments to our MAPseq data at the cubelet level, individual anterograde tracing experiments were normalised to injection volume, we then z-scored the Log_10_(mean Allen experiment summed voxel-wise projection density) within each cortical 3D LCM cubelet ROI location and compared this to the per mouse z-scored Log_10_(MAPseq barcode counts) for each LCM cubelet. To compare our MAPseq datasets to anterograde tracing experiments at the level of cortical areas, we calculated the mean z-scored Log_10_(projection density) for each cortical brain area across the 3 Allen experiments, and compared this to the Log_10_(mean projection density) to each area in our MAPseq dataset. Area projection density was calculated using “homogeneous across cubelet” area assignment approach (described in the Methods section: “Assignment of A1 projections to cortical brain areas”). The Pearson correlation coefficient and corresponding *p* value were calculated for either the summed MAPseq barcode counts for each cortical LCM sample or the mean projection density to each cortical area covered.

For comparison of projection strength to visual cortical areas based on injection location (Suppl. Fig.4b-d) using data from the Allen Mouse Brain Connectivity Atlas^33^, we included experiments where ≥95% of the injection of EGFP-expressing AAV was in auditory cortex (but not necessarily A1). Injections were divided into anterior or posterior based on injection location (anterior: 100149109, 120491896, 184159706, 182090318, 287223629; posterior: 116903230, 146858006, 569994739, 112881858). Some of the experiments were carried out in transgenic animals and featured Cre-dependent expression of EGFP (Cux2-IRES-Cre: 184159706, 569994739; Rorb-IRES2-Cre: 287223629; Rbp4-Cre_KL100: 182090318). Similar trends were observed when only including WT mice. Assignment of MAPseq neuron projection patterns to brain areas. Average projection strengths to visual areas were normalised within each animal. For analysis of relationship between A-P position of AC injection and projection strength to VC areas, a linear regression model was fit for all VC areas simultaneously using the glm function in the Statsmodels Python package. The model included Euclidean distance to each VC area as an additional fixed effect, in order to account for inter-areal distance. The resulting p-values were corrected for multiple comparisons using the Bonferroni approach.

#### Neuron projection assignment to broadly grouped brain regions

To calculate projection scores to broadly grouped brain areas (Fig. 1d), samples containing the cortex, thalamus, superior colliculus, inferior colliculus, and striatum were manually annotated during dissection. Counts were pooled across samples in brain region across annotated groups. Hierarchical clustering was performed using the clustermap function in the Python seaborn package^67^ with the canberra distance metric, with projection score determined as range normalised neuron barcode counts (subtracting minimum and dividing by range).

#### Assignment of A1 projections to cortical brain areas

Single neuron projection patterns to specific brain areas within the cortex were computed for IT neurons, identified as the neurons that did not project to either the thalamus or tectum. To assign barcode counts in cortical LCM cubelets to cortical areas, we employed an approach that assumed barcodes were distributed homogeneously across LCM cubelets and distributed barcode counts within a cubelet based on the fraction of the cubelet registered to each area. To calculate projection density to each area, we then summed the assigned neuron barcode counts for each area and divide by total area volume sampled. To this end, we constructed a matrix ***A*** containing samples and the volumes of brain areas contained within each sample, excluding areas that made up <10% of the cubelet. Since we were only interested in axonal projection patterns, we removed samples in the neuron barcode matrix that contained majority A1. We then computed matrix (*F*) containing proportion of each brain area as a fraction of the total volume of each LCM cubelet:

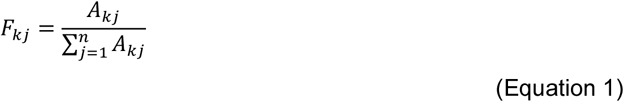

Where *F_kj_* is the element in the fraction matrix ***F*** at the *k*th cubelet and *j*t**h** area, *A_kj_* is the element in the area volume matrix *A* for the *k*th cubelet and *j*th area, with *n* representing the number of areas in matrix ***A.*** We then generated a weighted fraction matrix (***W***) and computed the matrix (***P***) of projection strengths for each neuron barcode across brain areas sampled by matrix multiplication:

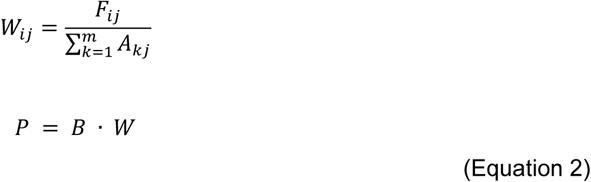

Where *W_ij_* is the element at the *i*th cubelet and *j*th area of the weighted fraction matrix (***W***), *F_ij_* is the element at the *i*th cubelet and *j*th area of the fraction matrix (***F***), *A_kj_* is the element at the *k*th cubelet and *j*th area of the areas in matrix (***A***), and *m* is the number of LCM cubelet samples. ***B*** denotes the matrix of barcode counts across LCM samples. To normalise projection densities, values were divided by their range so that each neuron’s maximum projection density equalled 1.

#### Comparison to other cortical area-assignment approaches

We compared the assignment of neuron projection patterns to cortical brain areas using the above approach to two other area assignment approaches: the “homogeneous across area” approach and the “area is main” approach. For the “homogeneous across area” approach, we assumed that neuron barcode counts were homogeneous across brain areas and therefore assigned neuron projection density to each area by multiple linear regression. Here, we computed the projection density to each brain area as the regression coefficient for the least squares solution of normalised barcode counts against area volume within each sample. We constrained the regression coefficient to be non-negative and used lasso regularisation with the *λ* parameter set to 1. As with the “homogeneous across cubelet” approach, we then normalised projection densities by dividing by the range.

For the “area is main” approach, we assumed that barcode counts in each cubelet derived from the brain area that made up the majority of that cubelet. This approach uses the same method as the “homogeneous across cubelet” approach, all non-majority area volumes in matrix **A** were set to zero.

We compared Pearson correlations between mice using different area assignment approaches and observed, on average, a higher correlation between mice for the “homogeneous across cubelet” approach (mean Pearson’s *R* = 0.87, 0.82, 0.83 for “homogeneous across cubelet”, “homogeneous across area”, and “area is main approaches”; Suppl. Fig. 2d-f).

### Analysis of single A1 neuron co-projection patterns

To analyse co-projection patterns of A1 neurons to different pairs of areas, we generated projection strengths of individual neurons to different brain areas using the “homogeneous across cubelet” approach.

#### Conditional probability of co-projection

To examine how co-projection patterns of neurons vary given they already target a specific cortical visual area, we calculated the conditional probability *P(target|VC area)* of A1 projection patterns:

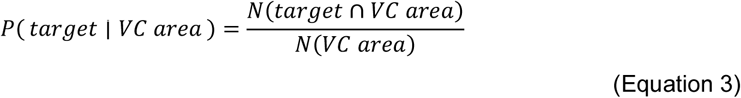

Where *P(target|VC area)* is the probability of projecting to a specific target area given that a neuron projects to a particular visual area, *N*(*target* ∩ *VC area*) is the number of neurons that project to both the target area the specified visual area, *N(VC area)* is the total number of neurons that project to that VC area.

We calculated conditional probabilities for all cortical areas present across all three mice in our dataset. To assess the similarity of cortical conditional projection probability patterns between different VC areas, we performed a cosine similarity analysis of conditional probability vectors using the cosine_similarity function in scikit-learn Python package.

#### Over- and under-represented motif analysis

Over- and under-represented co-projection motifs were identified by comparing the number of neurons co-projecting to a given pair of targets (‘*observed’*) with the number of co-projections expected by chance, based on the joint probability *P(*AB) of observing axonal co-projections to those areas (‘*expected’*).

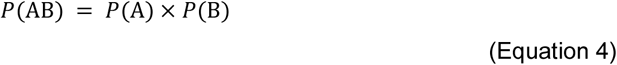

Where *P(AB)* is the probability of innervating both areas A and B together, *P(*A) and *P(*B) are the probability of innervating each area individually. ‘*Expected’* was calculated by multiplying the total number of neurons in the dataset with *P(*AB). We then computed the effect size as log_2_(*observed/expected*). Effect sizes were compared against the null shuffle population to assess statistical significance (see below).

#### Null shuffle population comparisons

Since the LCM cubelets contain mixtures of different brain areas, neurons within a cubelet are biased appear as if they co-project between pairs of areas present within the same LCM cubelet. To distinguish genuine neuron co-projection patterns from these within-cubelet biases, we generated null shuffle populations that preserve the same within-cubelet structure. Simulation of variation in barcode labelling efficiency in MAPseq can lead to artificial over-representation of projection motifs (Suppl. Fig. 3c). To generate the shuffle, we binarised cubelet barcode count matrices and then applied the curveball algorithm ^68^ to shuffle binarised barcode counts across neurons while preserving the total number of cubelet targets per neuron. This procedure thus maintained both the total number of neurons targeting each LCM cubelet and the total number of cubelet targets per neuron, thereby preserving neuron-specific differences in barcode labelling efficiency as well as the marginal probabilities of targeting cubelets. Using this approach, we performed 100,000 shuffles of neuron barcodes within LCM samples for each mouse. For every shuffle we reassigned projections to cortical brain areas using the ‘‘homogenous across cubelet’’ approach and then performed the corresponding conditional probability and co-projection motif analysis to generate a null population for each analysis type. Since the null distribution was approximately normal, we calculated two-tailed *p-*values by fitting a normal distribution to the shuffled data and computing the probability of observing a value at least as extreme as the effect observed in our data. *P-*values were then corrected using the Bonferroni method.

For visualisation of effect size in the conditional probability bubble plot (Fig. 2j), we plotted log_2_(observed conditional probability/mean shuffled conditional probability). For the co-projection motif volcano plot (Fig. 2l-k), we plotted the log_2_ fold change between the observed/expected co-projection motifs in the data and the corresponding observed/expected values from the shuffled data. We chose log fold difference rather than a simple log ratio of co-projecting neuron counts between observed and shuffled because shuffling preserves the marginal projection probabilities of individual cubelets, but not the tendency of the same neurons to target the same area across multiple cubelets. Consequently, the marginal probabilities after area assignment differ slightly between the observed and shuffled populations, making log fold difference of (observed/expected) in the observed data versus (observed/expected) in the shuffled data a more appropriate metric for effect size.

#### Simulation of barcode labelling efficiency

To examine how variation in barcode labelling efficiency can distort co-projection statistics in MAPseq datasets, we simulated detection of co-projection patterns from 100,000 neurons to 10 different brain areas under two efficiency scenarios: uniform or variable barcode labelling. Each neuron was given an independent probability of 0.2 for projecting to any one area. In the uniform labelling efficiency condition, every projection was detected with a probability of 0.5. In the variable labelling efficiency scenario, each neuron’s detection efficiency was an independent draw between 0 and 1. We simulated each neuron’s observed projection pattern by drawing a random probability between 0 and 1 for each neuron-target combination and subsequently recorded a projection to that target area if the random probability value was less than the efficiency-adjusted projection probability. This yielded observed neuron projection matrices used for subsequent co-projection analyses. For each area combination, we calculated expected co-projection values as described above for over- and under-represented motif analysis, based on the joint probability of observing co-projections calculated from total number of neurons projecting to each individual target area.

### Analysis of spatial relationships of A1 cortico-cortical projections in MAPseq dataset

#### Exponential decay with Euclidean distance

To calculate the relationship between projection frequency and projection strength (normalised barcode counts per mm^3^ area) with distance from A1, we calculated Euclidean distances from area centroids in Allen CCF space. For A1 projections to cortical areas covered by all three mice in our dataset, we fitted the change in projection frequency and projection strength with Euclidean distance using the exponential decay model:

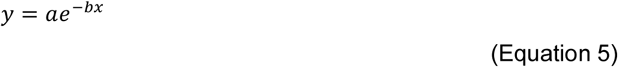

Where *y* is the dependent variable (projection frequency or projection strength), *a* is the predicted value of *y* at 0 distance, *b* is the decay constant (µm^−1^), and *x* is the Euclidean distance between A1 and area centroids (µm).

The exponential decay model was fitted using the optimize.curve_fit function in the SciPy Python package. To assess goodness of fit, we used leave-one-out cross-validation to predict the projection frequency and strength for each sample and computed the *p*-values of the Pearson correlation coefficients between cross-validated predictions and observed values.

#### Topographic mapping of MAPseq projection patterns

To examine the relationship between A-P soma position and A-P projection patterns at the cubelet level, we computed the A-P coordinates of A1 source and VC target cubelet centroids, normalised such that the most posterior position in VC corresponded to 0 for both source and target. VC cubelets containing >10% A1 (1 cubelet) were excluded to minimize confounding from local A1 projections. For each cubelet A-P position containing more than two projecting neurons, we calculated the mean A-P position of the corresponding somata. To extend this analysis to the level of VC areas, we calculated the mean A-P soma position of neurons projecting to each VC area (assigned using the “homogeneous across cubelet” approach) for each mouse individually.

To assess the relationship between targeting frequency of individual areas and A-P position, we generated binary single neuron projection matrices to each VC area for each soma A-P position. Cubelets containing fewer than five neurons were excluded. Soma A-P position was treated as a continuous variable, and single neuron projections to individual VC area as a binary variable. To test whether projection probability varied with A-P position, logistic regression models were then fitted for all VC areas simultaneously using the Statsmodels Python package. Here, the logistic model relates projection probability to A-P soma position, allowing baseline projection probability and slope to differ between VC areas, and includes Euclidean distance to the target VC area and mouse as additional covariates. Euclidean distance was calculated as the distance between the centroids of each VC area and the A1 soma cubelet at each A-P position. This model therefore accounts for within-mouse variability in baseline projection probabilities and the effect of inter-areal distance by setting them as fixed effects. The resulting p-values were corrected for multiple comparisons using the Bonferroni approach.

#### Additional resources for MAPseq data visualisation

Brainrender was used to generate 3D videos and images of MAPseq datasets^69^. To visualise data on cortical flatmaps, we implemented the ccf_streamlines Python package ^70^, available from https://github.com/AllenInstitute/ccf_streamlines.

### Verification of injection sites after functional imaging

Mice were euthanized with a dose of pentobarbital (80 mg/kg) and transcardially perfused with 4% paraformaldehyde. Brains were extracted and post-fixed overnight in 4% paraformaldehyde. Brains were embedded in 5% agarose and then sliced and imaged with a custom-built serial-section two photon imaging system ^71^, controlled by Scanimage (Vidrio Techonolgies) using BakingTray (https://github.com/SWC-Advanced-Microscopy/BakingTray). Brains were imaged at 920 nm, with slices imaged every 10 μm along the antero-posterior axis, with a resolution of 2 μm per pixel. In some cases, slices were collected afterwards and imaged at higher resolution on a confocal microscope (Nikon CSU-W1).

### Intrinsic signal imaging

Intrinsic imaging was performed under isoflurane anaesthesia. Mice were injected with acepromazine (3 mg/kg), ∼10 min prior to imaging. Isoflurane was administered at a relatively low concentration (∼0.7%) during imaging. The surface of the brain was illuminated at 700 nm with 2 LEDs (Thorlabs M700L4) coupled to light-guides (Thorlabs). Images were acquired with a CMOS camera (acA2040-120cm, Basler) through a custom-built microscope, using a Nikon (AF Nikkor 50mm f/1.8D) objective. The objective was focused at −500 μm from the brain surface during imaging. Bonsai (Lopes et al., 2015) was used for data acquisition and synchronization. Visual stimuli were generated in MATLAB, and presented on an LCD screen (U2415b, Dell), located 20 cm from the mouse’s head, covering 105° in azimuth and 76° in elevation. A checkered bar drifting horizontally or vertically across the screen was presented, at a speed of 15°/s. The bar drifted 10 times in each of the four cardinal directions. Retinotopic maps were computed by taking the phase of the response (change in tissue reflectance) recorded at each pixel, relative to the phase of the drifting bar^72^. Maps obtained from stimuli drifting in opposite directions were subtracted to correct for haemodynamic response delays.

### Two-photon imaging

Two-photon imaging was performed 3-10 weeks after viral injection, through a 16x/0.8-NA objective (Nikon) using a custom-built resonant scanning two-photon microscope (INSS), controlled by Scanimage (Vidrio Technologies^73^). A Ti:Sapphire femtosecond-pulsed laser (Chameleon Ultra II, Coherent) was used to stimulate the fluorophores at 930 nm (for imaging jGCaMP7b only) or 1010 nm (for imaging jGCaMP7b and jRGECO1a simultaneously). Power post objective was between 60 and 100 mW, depending on expression and FOV quality. Field of view sizes were between 270×270 µm and 520×520 µm. Typically, 4 optical planes were acquired quasi-simultaneously at 6.7 or 6 Hz, using a piezo actuator (Physik Instrumente) attached to the objective to rapidly switch between imaging planes. Laser power was adjusted with imaging depth, to obtain roughly similar fluorescence levels across planes. Axons were imaged between 20 and 100 µm below the pia, VC-neuron somata were imaged 50-150 µm below the pia.

### Auditory and visual stimuli

Stimuli were generated in MATLAB (2018a, MathWorks) using custom code, and presented using Psychotoolbox-3^74^. Sound stimuli were digital-to-analog converted using a soundcard (ASUS Xonar X7 or PEXSOUND7CH), with a 192 kHz sampling rate.

#### Stimulus location experiments

For stimulus location experiments, auditory and visual stimuli were presented using an array of concentric speakers and LEDs, organised in a 3-dimensional custom-built structure with a 20 cm radius (see Figure 3l). The speakers and LEDs were all identical (speakers: VISATON, K 28 WPC BL, LEDs: 2.4 lux, 4.3°, OSRAM SmartLED, LWL283-Q1R2-3K8L-1). Further details of the stimulation apparatus can be found at https://github.com/Iacaruso-lab/Coliseum. A 1.5 cm diameter white acrylic diffuser was placed in front of each LED, to increase the size of the visual stimulus. All SPLs reported for these speakers are in presence of these diffusers. Sound calibration was done using a free-field microphone (GRAS 46BE) coupled to an audio analyser (APx517). The microphone was positioned 0.5 cm from the diffuser for calibration. We verified all speakers produced sound at the same intensity (± 3 dB, inter-speaker variability was in same range than variability due to slightly different microphone positioning relative to speaker).

The sound was composed of a series of short bursts of bandpass-filtered white noise (4-24 kHz), presented for 500 ms (4 bursts total, 50 ms per burst, 75 ms between bursts), with a variable inter-trial interval (1.5-2.5 seconds). The sounds were presented at slightly varying SPLs, centred around 65 dB (± 5 dB). Sounds were presented at 3 different elevations (−36°, 0° and 36°), and either 7 or 13 different azimuths, spanning the ipsi-and contralateral hemifields (−108° to 108°, with either 18° or 36° between speakers). Sound source locations were randomised across trials.

Visual stimuli used for retinotopic mapping consisted of a flashing white LED, presented in one location at a time. The LED’s flashing pattern emulated the noise bursts in the sound stimulus (i.e 4 flashes, 50 ms duration each,75 ms between flashes, total length of 500 ms). Visual stimuli were presented at 3 or 5 different elevations (spanning −36° to +36°, with either 18° or 36° between LEDs), and 4 or 7 different azimuths, spanning 0° to 108° on the contralateral side (with either 18° or 36° between LEDs).

Visual and auditory stimuli were presented either interleaved with each other in random order, or in separate blocks (i.e. first a visual only block and then an auditory block), and were never presented at the same time. No other light was present during the recording.

#### Pure tone experiments

Pure tone stimuli were presented through an amplifier (Avisoft Bioacoustics) and played through a free-field electrostatic speaker (Vifa, Avisoft Bioacoustics). The speaker was coupled to a cone, padded with cotton to minimise reverberations. The tip of this cone was placed ∼ 1 cm from the entrance to the mouse’s ear canal, on the contralateral side. Stimuli consisted of pure tones with different frequencies (2-64 kHz, ½ octave steps), which were presented at 3 different sound pressure levels each (40-, 50- and 60-dB SPL). Tones were presented in random order, for 200 ms each, with 2 seconds between trials. An LCD screen (U2415b, Dell) was on at all times during these experiments, displaying a gray screen.

### Behavioural monitoring

During all recordings, videos of the face and the body were recorded at 15 Hz through two separate cameras. The recording setup was illuminated with an infrared light. The face was recorded through an infrared camera (DMK 22BUC03), coupled to a 50 mm lens (computar). The body was recorded through an additional infrared camera (Logitech). Additionally, locomotion speed was monitored using a rotary encoder (Kuebler), located at the wheel’s axle. Acquisition of behavioural data and synchronisation with two-photon imaging and stimulus presentation was done using Bonsai.

### Imaging data analysis

Data analysis was carried out using custom code in Python and MATLAB, except were indicated.

#### Quantification of injection spread and location

Histology images obtained from serial two-photon tomography were assembled using StitchIt (https://github.com/SWC-Advanced-Microscopy/StitchIt). Brain slices were aligned to the Allen Mouse Brain Atlas^31^ using brainreg^75^ (https://github.com/brainglobe/brainreg), and the area of expression was then manually traced using brainreg-segment (https://github.com/brainglobe/brainglobe-segmentation). The segmented region was then used to calculate the percentage of expression in each area, and to visualise expression locations in the aligned cortical flatmap (Suppl. Figure 5b). Injection centroids were taken as the centre of the flatmap-projected segmented region.

#### Retinotopic map alignment

Intrinsic imaging maps for azimuth and elevation were obtained for each animal as explained above. A template outline of higher visual areas was manually aligned to each animal’s retinotopic maps, using known features of area boundaries such as the reversal of retinotopy in the borders of V1-LM, V1-RL, V1-PM, AL-LM, and AM-PM. ROIs (regions-of-interest, boutons or somata) were assigned to visual areas based on the position of the aligned area boundaries. In cases where fields of view were on the border between two areas, ROIs were assigned to the area they are located in. For session-wise analysis, all ROIs in a given FOV were assigned to the area that contained most of the ROIs. For cross-animal visualisations and analysis, the area outlines were then registered across animals using affine transformation, implemented using the bUnwarpJ package in ImageJ (https://imagej.net/plugins/bunwarpj/;^76^). The obtained transformation matrices were then used to register ROI locations across animals.

#### Two-photon imaging

Imaging movies were registered, and regions-of-interest (ROIs) were extracted using the python implementation of the suite2p toolbox^77^, https://github.com/MouseLand/suite2p). Frames in which registration motion was large (motion > 5 pixels) were excluded from further analysis. For somatic data, cells were manually sorted based on visual inspection of the registered frames. For axonal data, the threshold for adding new pixels to ROIs during the “ROI extension” phase of ROI extraction in suite2p was increased, which largely prevented bouton ROIs from also incorporating axonal segments and other nearby boutons. Then, putative boutons were automatically detected in the average registered image using a blob detection algorithm based on a difference of Gaussians approach (python skimage library, feature.blob_dog function). The standard deviation of the Gaussian kernels was empirically set such that individual boutons were recognised as blobs, while larger axonal fragments and occasional multi-bouton clusters were not. Only suite2p axonal ROIs that co-localised with these blobs were considered to be axonal boutons and included in further analysis. In somatic ROIs, fluorescence was neuropil corrected, using (F = F_raw_ – 0.7*F_neu_), were F_raw_ was the fluorescence inside the ROI and F_neu_ was the estimate of the neuropil fluorescence. In all ROIs, fluorescence was normalised to the baseline as ΔF/F = (F-F_0_)/F_0_, where F_0_ was calculated as the 30^th^ percentile of the fluorescence over the entire recording.

#### Generalised linear model (GLM)

Before further analysis, a generalised linear model was fit to each boutons’s fluorescence across time, using a set of motor and stimulus predictor variables (Suppl. Fig.7). Neural activity was baseline subtracted, and fluorescence intensity was binned into 100 equally spaced bins. Since the sampling rate of two-photon imaging is relatively low (6-6.7 Hz in these data), data was not further binned in time. Motor predictor variables comprised the locomotion trace, as well as the top 30 non-laser-flicker (see above) principal components of the mouse face motion. Each motor predictor variable was grouped into 10 bins for the model. Stimulus variables comprised one separate variable for each stimulus ID (i.e. each stimulus location or frequency-sound intensity combination). In sessions with visual and auditory stimuli, each stimulus location had one auditory and one visual variable. All these variables had a value of 1 when the stimulus they encoded was present, and 0 at all other times. In addition, a variable encoding time during the trial was added.

The model implemented was taken from Hardcastle et al.^78^ and Yoo et al.^79^ and adapted for Gaussian-distributed data. The model estimates the neural activity *f* of a bouton/neuron during time t as:

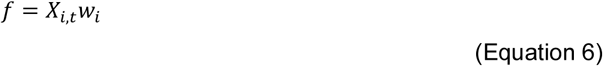

Were *X_i,t_* is the value of a predictor variable *i* at a given time point *t* and *w_i_* contain the weights for each variable *i.* Models were fit using the fminunc function in MATLAB, using five-fold cross-validation, and additional regularization for the smoothness of parameters in a continuous variable, and a lasso regularization for parameters in discrete variables. Model performance was assessed with cross-validated variance explained (cVE), calculated as:

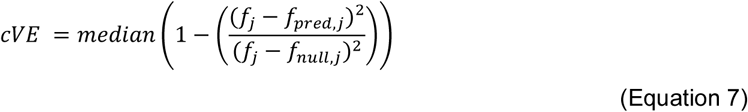

where *f* is the neural activity of the bouton/neuron in one of the *j* folds, *f*_pred,j_ is the neural activity predicted from the model using the fit weights, and *f*_null,j_ is the mean of *f*. This measure is similar to R^2^ in simple linear regression, and quantifies how close the predicted trace *f_pred_* is to the measured trace *f*, relative to a null prediction. A model with a set of predictors was considered significant if the variance explained by those predictors was significantly higher than zero across folds, using a one-sample Wilcoxon signed rank test.

#### Motor component subtraction

Subtraction of the motor component was done by subtracting the fluorescence trace predicted based on the fitted motor variables from the measured fluorescence trace. To evaluate whether this subtraction removed tuning to motor variables, each motor variable was grouped into 10 or 15 equally spaced bins according to their magnitude. Tuning curves were computed for each bouton by averaging the fluorescence within each of these behavioural bins. Fluorescence was z-scored before averaging by behaviour bins, and the resulting tuning curves were sometimes normalised between 0 and 1 for visualisation. To evaluate tuning to motor variables, the same approach as Christensen and Pillow^80^ was used. Boutons were considered to be tuned to a behavioural variable (locomotion, facial motion) if the variance of their fluorescence across behavioural bins was significantly higher than the variance of the fluorescence shuffled across behavioural bins (Levene’s t-test of variance).

#### Stimulus tuning analysis

For analysis of neural responses to stimuli, the motor component predicted from the motor linear model described above was subtracted from each roi’s fluorescence trace. The motor-subtracted traces were aligned to stimulus presentation, and the response was calculated as *R* = *F_resp_* – *F_base_*, where *F_resp_* is the average ΔF/F over the 1s following stimulus onset and *F_base_* is the average ΔF/F over the 0.5-1 seconds (depending on inter-trial interval) preceding the stimulus. This *R* value was averaged across repeated presentations of the same stimulus to obtain frequency-response areas (FRAs) or location response areas (LRAs)

Neurons were classified as responsive if the *F_resp_* across trials for the stimulus type evoking the strongest mean *F_resp_* (stimulus location or frequency-SPL combination) was significantly higher than the *F_base_* (one sided Wilcoxon signed-rank test) and the mean *F_resp_* for that stimulus exceeded 1 ΔF/F. Neurons were classified as azimuth or elevation-selective if they were responsive and exhibited a significant change in response with azimuth or elevation respectively, assessed by a two-way ANOVA (p <0.05). Similarly, neurons were classified as tone frequency-selective if they were responsive and presented a significant change in response with tone frequency, assessed using a two-way ANOVA. Only sessions with at least 10 selective boutons were included in analysis that treated sessions as units.

Analysis of location or frequency tuning was performed on trial-averaged responses. To assess tuning to sound azimuth, responses were averaged across stimulus elevations, and the resulting 1-D vector was fit with a 1-dimensional Gaussian curve. The peak of this Gaussian was taken as the bouton’s “best azimuth”. Only boutons with an *r*^2^ of Gaussian fit >0.6 were included in analysis of best azimuth. Results were qualitatively similar when defining best azimuth as the azimuth evoking the highest ΔF/F, without fitting a Gaussian. Azimuth tuning width was calculated as the full-width at half-max (FWHM) of the Gaussian fit to the azimuth tuning curve.

To calculate the best elevation for each bouton, responses across azimuths within the bouton’s preferred visual field portion (ipsilateral, centre or contralateral) were averaged. The elevation that evoked the strongest activity was taken as “best elevation”. Best visual azimuth and elevation in VC neurons were obtained by fitting a 2-dimensional Gaussian to their 2-dimensional receptive fields. For cases where the *r*^2^ of the fit was < 0.3, the azimuth and elevation evoking the highest average ΔF/F were taken instead.

To quantify tuning to tone frequency, responses across all sound intensities for each frequency were averaged. The resulting frequency tuning curve was fit with a spline, and the highest peak was taken as “best frequency”. Frequency tuning width was calculated as the FWHM of the spline fit. Boutons were classified as “multi-peaked” if responses to a frequency at least one octave away from the best frequency were at least 75% as strong as the response to the best frequency. Only non-double peaked boutons were included in best frequency analysis. To calculate the sparsity index (SI) of pure tone responses, the following formula was used^81^:

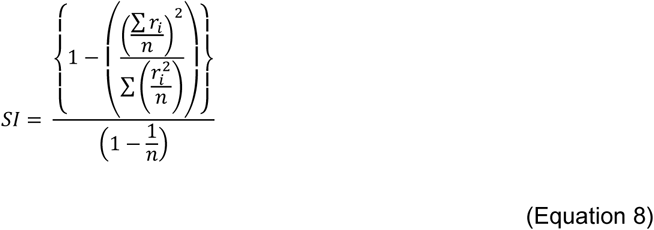

Where r_i_ denotes the average response of a given bouton to a given stimulus (i.e. a combination of tone frequency and intensity) and n is the number of different stimuli.

#### Signal correlations

Paired signal correlations between axons were calculated as the Pearson correlation between their trial-averaged stimulus responses (i.e. their LRAs or FRAs). For signal correlations involving azimuth or elevation representations, responses were averaged across the non-relevant dimension in each case.

#### Grouping of boutons into axons

We used an interactive clustering procedure based on correlations in activity over time to group boutons belonging to the same axon, similar to the procedure described in Mazo et al^29^. For each session, a random pair of boutons with a Pearson correlation above a threshold of 0.4 was selected as the initial seed for a cluster. Additional boutons were then assigned to that cluster if their correlation with any bouton in the cluster exceeded the threshold, or they initiated their own cluster otherwise. For each cluster, the bouton with the highest average ΔF/F was selected to represent that axon. The threshold 0.4 was determined based on empirical correlations between boutons belonging to the same axon versus different axons.

#### Hierarchical bootstrap

A hierarchical bootstrap approach was used for analysis at the bouton/axon level, as described in refs^82^. For each bootstrap sample, mice were resampled with replacement, followed by resampling of experimental sessions and subsequently boutons/axons. A statistic of interest was then calculated for each bootstrap sample. This statistic was either the difference in means (for signal correlations) or the difference in medians (for frequency tuning). We then calculated the quantile *q* of the bootstrap distribution at 0, and calculated the two-sided bootstrap *p*-value as 2 × min{*q*, 1 − *q*}.

#### Linear mixed models

Linear mixed models (LMMs) were used for session-level analyses and implemented using the MATLAB function fitlme. LMMs were primarily used to assess correlations between variables or to test for differences across areas, in combination with a mixed effects ANOVA. In all models, the antero-posterior and medio-lateral positions of the injection site in each animal were included as fixed effects, while animal identity was included as a random effect. For example, when analysing best sound elevation, the model included an intercept, visual area as a categorical fixed effect, injection site coordinates (anteroposterior and mediolateral) as covariates, and a random intercept for animal.

### Facial motion analysis

Facial motion energy was calculated as the frame-to-frame change in luminance for each pixel in the face video, averaged across all pixels. In some cases, motion energy traces were filtered to remove oscillations caused by laser intensity changes with imaging depth (Butterworth stopband filter centred at 3 Hz). Facial principal components (PCs) were obtained via singular value decomposition, and projected back on the time axis, using the algorithm implemented in the facemap toolbox^83^, https://github.com/MouseLand/facemap). Some facial motion PCs were dominated with the laser flicker, while others were unaffected. PCs with a strong 3 Hz spectral component were excluded from further analysis. Locomotion speed was recorded via a rotary encoder attached to the wheel axle. All behavioural variables were downsampled to the two-photon acquisition rate (6-6.7 Hz) prior to analysis.

#### Stimulus decoding from facial motion

A linear support vector machine (SVM) classifier was used to decode stimulus type (sound azimuth, elevation, tone frequency or tone intensity) using the first 100 PCs of facial motion. Motion in each PC was averaged over the response period and z-scored before classification. Trials were pooled across non-relevant dimensions (e.g. for decoding stimulus azimuth, trials were pooled across elevations). The classifier was trained and tested on held-out data using 10-fold cross validation. For each session, the classifier was run 10 times, subsampling 50 trials of each stimulus type at random without replacement per iteration. Average accuracy across stimulus types was then computed across iterations.

### Other statistical analysis

After testing for differences in stimulus tuning across areas using the methods described above, pairwise comparisons between areas were performed using the Mann–Whitney U test. To control the false discovery rate due to multiple testing, the Benjamini–Hochberg procedure was applied to the resulting *p*-values.

### Data availability

Preprocessed data required to reproduce the analyses will be deposited and made publicly available prior to publication. Both preprocessed and raw data are available from Florencia Iacaruso (florencia.iacaruso{at}crick.ac.uk) upon request.

### Code availability

The open-source designs of the speakers and LED device and its associated circuits is available on GitHub (https://github.com/Iacaruso-lab/Coliseum). The code for pre-processing and registration of raw MAPseq datasets is available at https://github.com/znamlab/MAPseq_processing. Matlab and python code for analysing the *in vivo* data and for reproducing the MAPseq data analyses is available on GitHub (https://archive.softwareheritage.org/swh:1:snp:bfb8c2237701bb3d01b702e1c8b49ed17afd3919;origin=https://github.com/Iacaruso-lab/AC-VC-paper).

## Acknowledgments

We thank Johannes Kohl and members of the Neuronal Circuits and Behaviour Lab and the Specification and Function of Neural Circuits Lab for discussions and comments on the manuscript. We thank the Biological Research and Surgical Services Facilities at the Francis Crick Institute for animal care and technical assistance, as well as Crick Light Microscopy, Experimental Histopathology, Genomics, Making Lab, Mechanical Workshop, and Vector Core. This study received support from the Francis Crick Institute, core funding (M.F.I: 10746, CC2118,P.Z.CC2108). The Francis Crick Institute receives its funding and the Francis Crick Institute which receives its core funding from Cancer Research UK, the UK Medical Research Council, and the Wellcome Trust. The work was also supported by the Engineering and Physical Sciences Research Council (BBSRC, award ref. EP/X020924/1, M.F.I)

## Author contributions

A.E.W., B.T.B., P.Z. and M.F.I. conceptualised the study. A.E.W., B.T.B. & A.V. performed experiments, A.E.W., B.T.B., A.V. & E.M. performed preliminary analysis. A.E.W. & B.T.B. performed formal analysis. P.Z. & M.F.I. acquired funding. P.Z. & M.F.I. supervised the project. A.E.W., B.T.B., P.Z. & M.F.I. wrote the paper.

## Supplementary Methods

### Optimisation of second strand synthesis approach for MAPseq library preparation

We tested whether targeted primers for 2^nd^ strand synthesis improve the efficiency compared to RNA template fragments. 100ng of spike-in RNA was used for each 20μl reverse transcription reaction. Reverse transcription was performed using SuperScript IV (Thermo Fisher Scientific, #18090050) with the gene-specific reverse transcription primer (Optimization_RT_001) using conditions described previously^30^. Here, the primer annealing step was carried out at 70 °C for 10 min, followed by reverse transcription at 55 °C for 10 mins, followed by enzyme deactivation at 80°C for 10 min. The resulting first-strand cDNA was divided into two aliquots for either: (a) targeted primer based or (b) RNA template as primer based second strand synthesis. For (a), 0.5 μl RNAse H (New England Biolabs, #M0297L) was added to 10μl of 1^st^ strand reaction and incubated for 20mins at 37°C. Second strand synthesis was then carried out by adding 0.75 μl primer (Optimization_PCR_002), 0.2 μl AccuPrime Pfx (Thermo Fisher Scientific, #12344032), 2.5 μl 10x buffer, and water to a final volume of 25μl. The sample was heated to 95 °C for 2.25 secs, then 68 °C for 2 min, and placed on ice. For (b), 2^nd^ strand synthesis was performed as previously described^30^. Per 10 μl first-strand cDNA, 6.68 μl water, 4.68 μl second strand buffer, 0.63 μl 10mM dNTP (Thermo Scientific, #10319879), 0.21 μl RNase H (New England Biolabs, #M0297L), 0.84 μl DNA Polymerase I (TaKaRa #2130A), 0.21 μl *E. coli* DNA ligase (New England Biolabs #M0205S) were added and incubated at 16 °C for 2 hours, then placed on ice. Subsequently, 0.84 μl T4 DNA polymerase (New England Biolabs #M0203S) was added and incubated at 16 °C for 10 mins, then placed on ice.

Both (a) and (b) second strand synthesis reactions were purified with a 2x bead clean-up using KAPA Pure Beads (Roche, #KK8001), washed twice with 80% ethanol, and eluted in 10 μl water. 8 μl of the second stranded cDNA eluate was exonuclease treated by adding 1μl Thermolabile Exonuclease I (New England Biolabs, M0568L) and 1 μl of buffer r3.1, and incubated in a thermocycler for 60 mins at 37 °C followed by 20 min at 80 °C. ExoI-treated cDNA was then amplified by nested PCR. The first round of PCR was performed for 12 cycles using AccuPrime Pfx with primers Optimization_PCR_002 and Optimization_PCR_003 and 10 μl of cDNA. 10μl Thermolabile Exonuclease I was then added to the 50 μl PCR reaction, incubated at 37 °C for 30 min and heat inactivated at 80 °C for 20 min. The second PCR reaction was performed using primers Optimization_PCR_004 and Optimization_PCR_005 (see primer table for oligo sequences) for 5 cycles on 7.5 μl of the PCR product. 20 μl of the resulting PCR product was purified using Wizard SV Gel and PCR Clean-Up System (Promega, #A9281) and quantified using an Agilent 4200 TapeStation. We compared the PCR product concentrations generated from cDNA from the original or targeted second strand synthesis approaches using a paired *t*-test.

## Supplementary figures

**Supplementary Figure 1.**
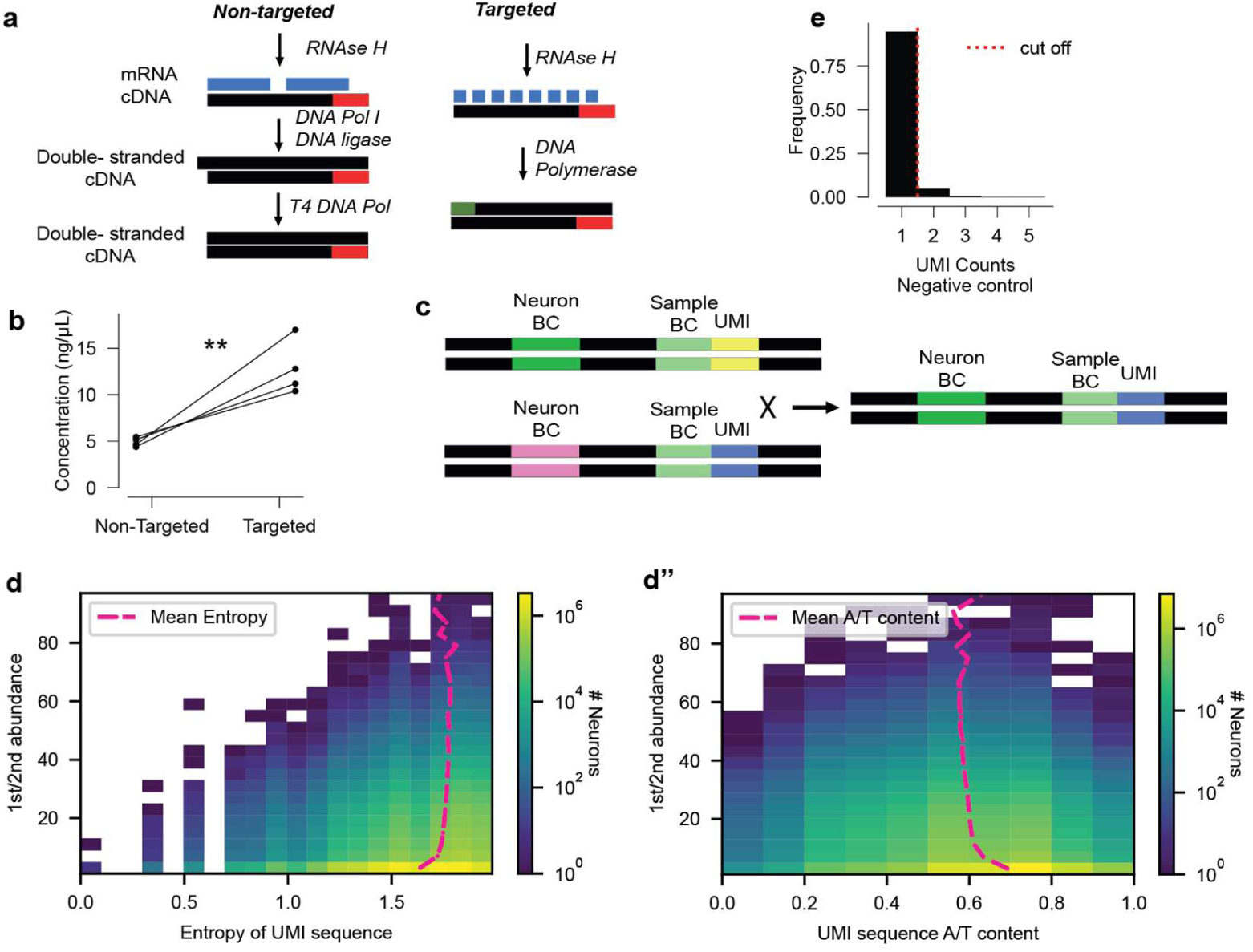
Optimisation of MAPseq Next Generation Sequencing (NGS) library preparation and identification of template switching events. **a,** Summary of different second strand synthesis approaches compared in optimisation of MAPseq sequencing library preparation. In the non-targeted approach, the nicked mRNA is used as primers for second strand synthesis, while in the targeted approach, RNA is degraded and a targeted primer anneals to the single stranded DNA and used for second strand synthesis. **b,** Comparison of non-targeted and targeted second strand synthesis approaches in MAPseq library preparation shows greater yield of final PCR product using the targeted approach. N = 4, *p*=0.017, paired two-tailed *t-*test. **c,** Diagram illustrating template switching during PCR amplification in MAPseq library preparation. **d,** 2D histogram showing the unique molecular identifier (UMI) count for neuron barcodes with the same UMI, where 1^st^_max_ is the UMI count for the neuron-barcode/UMI combination that is most abundant, and 2^nd^_max_ is UMI count for the neuron-barcode/UMI combination that is second most abundant. At 1^st^_max_/2^nd^_max_ ratios of less than 10 we observe a decrease in entropy, as well as **d’,** an increase in AT content in the UMI sequence, suggesting that shared UMIs with neuron barcodes with similar UMI counts are resultant of sequencing artifacts or biases in random nucleotide synthesis during oligo synthesis. Ratios with 1^st^_max_/2^nd^_max_ ≥10 abundance were therefore considered template switching events and only the most abundant neuron-barcode/UMI combination kept, while below this ratio all reads were discarded. **e,** UMI count distribution for negative control (olfactory bulb) LCM samples in MAPseq dataset. Setting a UMI count threshold of ≥2 removes 95% of barcodes from negative control.

**Supplementary Figure 2.**
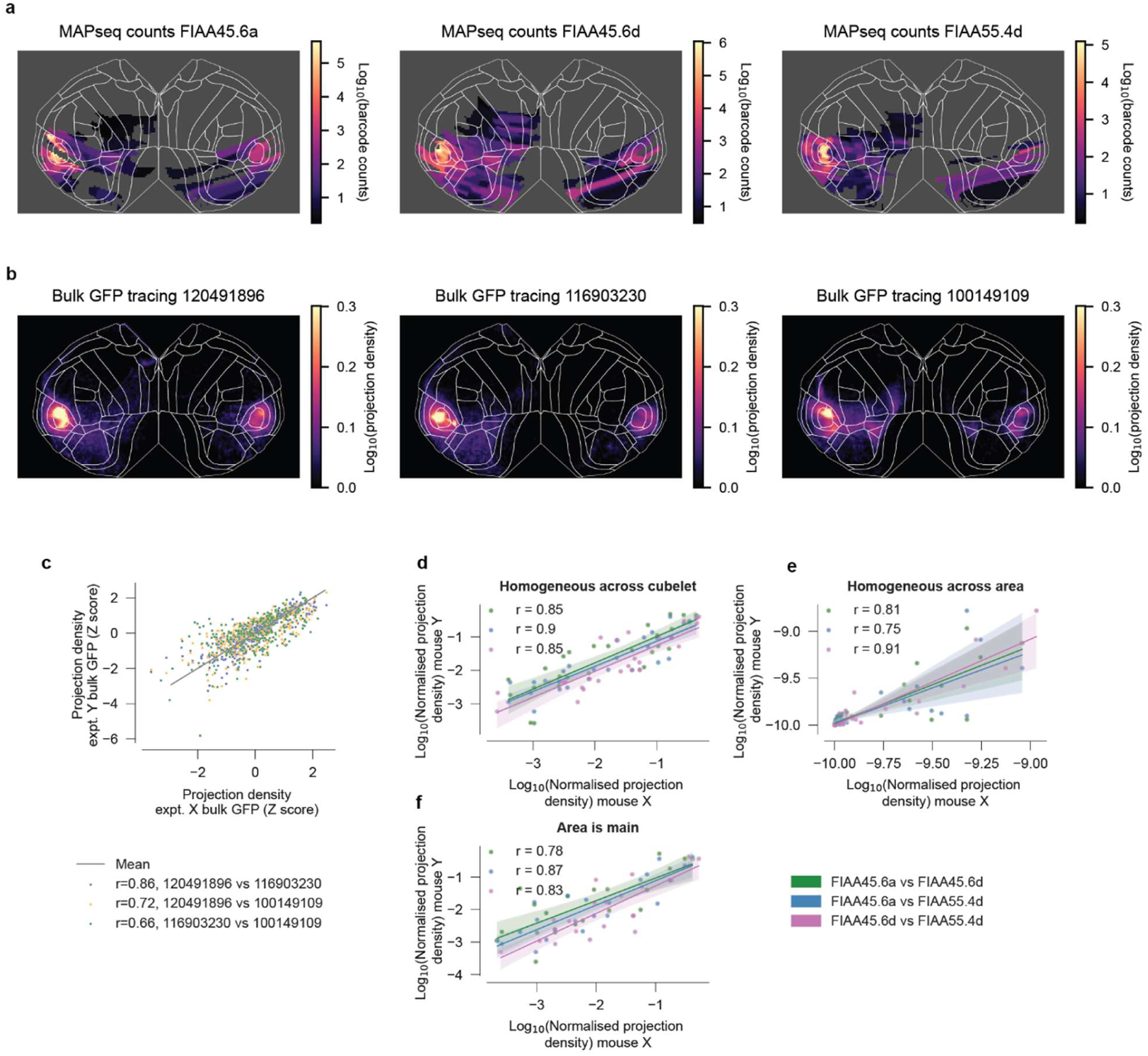
MAPseq comparison to anterograde tracing datasets, and barcode area assignment approaches. **a,** Cortical flatmaps showing distribution of bulk MAPseq barcode counts across cubelets for individual mice. **b,** Flatmaps of summed projection strength of bulk anterograde GFP tracing for three different experiments from the Allen Connectivity Atlas. **c,** Z-scored bulk projection density for individual Allen Connectivity Atlas anterograde tracing experiments plotted against each other. **d,** Comparison of mean area-specific normalised projection density between mice using the “homogeneous across cubelet” (where barcodes are assumed to be homogeneously distributed across cubelets) **e, “**homogeneous across area,” in which barcode distribution is assumed to be homogeneous across brain areas, and **f, “**area is main” where barcode counts assigned to brain regions with largest representation within each cubelet (see methods).

**Supplementary Figure 3.**
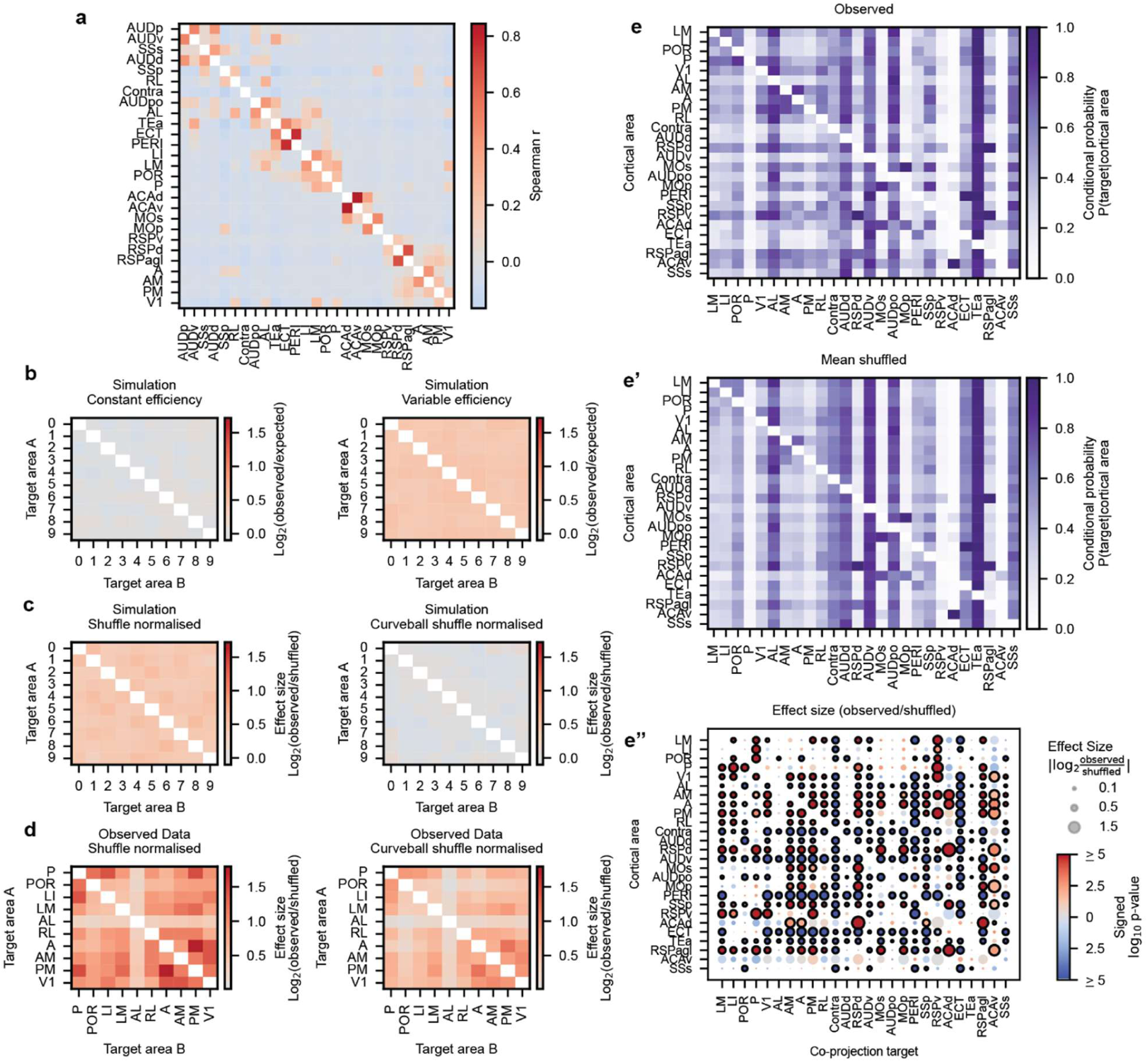
Identifying overrepresented co-projection patterns while controlling for intrinsic cubelet biases and variation in labelling efficiency. **a,** Heatmap showing mean Spearman correlation coefficient between cortical brain areas in LCM samples across all 3 mice. In addition to biases in observed co-projections based on cubelet structure, axon projection labelling efficiency by MAPseq Sindbis virus can vary between neurons. Neurons with lower labelling efficiency would have fewer barcode counts in their target areas, which would lead to some projection targets being missed due to limits in sensitivity. **b,** To test whether differences in labelling efficiency affected observation of co-projection motifs, we simulated observed/expected number of co-projections with either equal efficiency of labelling (left) or variation in neuron labelling efficiency (right). In the simulation, 100,000 neurons were sampled with p=0.2 of targeting each of the 10 areas and given either a constant or variable labelling efficiency. Expected co-projections are the product of individual marginal probabilities (independence assumption). We observed that simulated differences in axon barcode labelling efficiency can lead to artificial over-representation of co-projection motifs. **c,** To correct for spurious over-represented co-projection motifs that arise when two target areas fall within the same cubelet, we compared observed data to the same dataset after shuffling neuron barcode identifies within cubelets. However, normalising by the log₂ fold change between observed and shuffled simulations revealed that simple random shuffling (left) does not correct for the overrepresentation bias due to variable efficiency. In contrast, shuffling using the curveball algorithm (right) which preserves individual neuron labelling efficiencies was able to correct for artificial overrepresentation due to variable labelling efficiencies in the simulation. **d,** As predicted by simulations, applying this same principle to our MAPseq dataset showed that correcting for both variable labelling efficiencies and shared areas within cubelets using the curveball shuffle approach (right) decreased co-projection motif effect sizes compared to the random shuffle approach (which only corrects for shared areas within cubelets, left). **e,** Heatmaps showing conditional probabilities of A1 neurons targeting cortical co-projection targets ‘Co-projection target’ given neurons already target a given cortical area (‘Cortical area’). **e,** shows actual data conditional probabilities. To determine statistical significance from chance, we used a shuffle population of 100,000 shuffles, maintaining differences in labelling efficiency and marginal probabilities (see Methods). For visualisation, the **e’** heatmap shows the mean of 100,000 shuffles, and **e’’,** shows the effect size (log2(observed/shuffled conditional probabilities) red and blue circle colours indicate increased and decreased effect sizes respectively, shading indicates strength of *p*-value (as indicated on the colour-bar), black outline indicates *p*≤0.05. *P*-values are calculated from the null shuffled population distribution after Bonferroni correction.

**Supplementary Figure 4.**
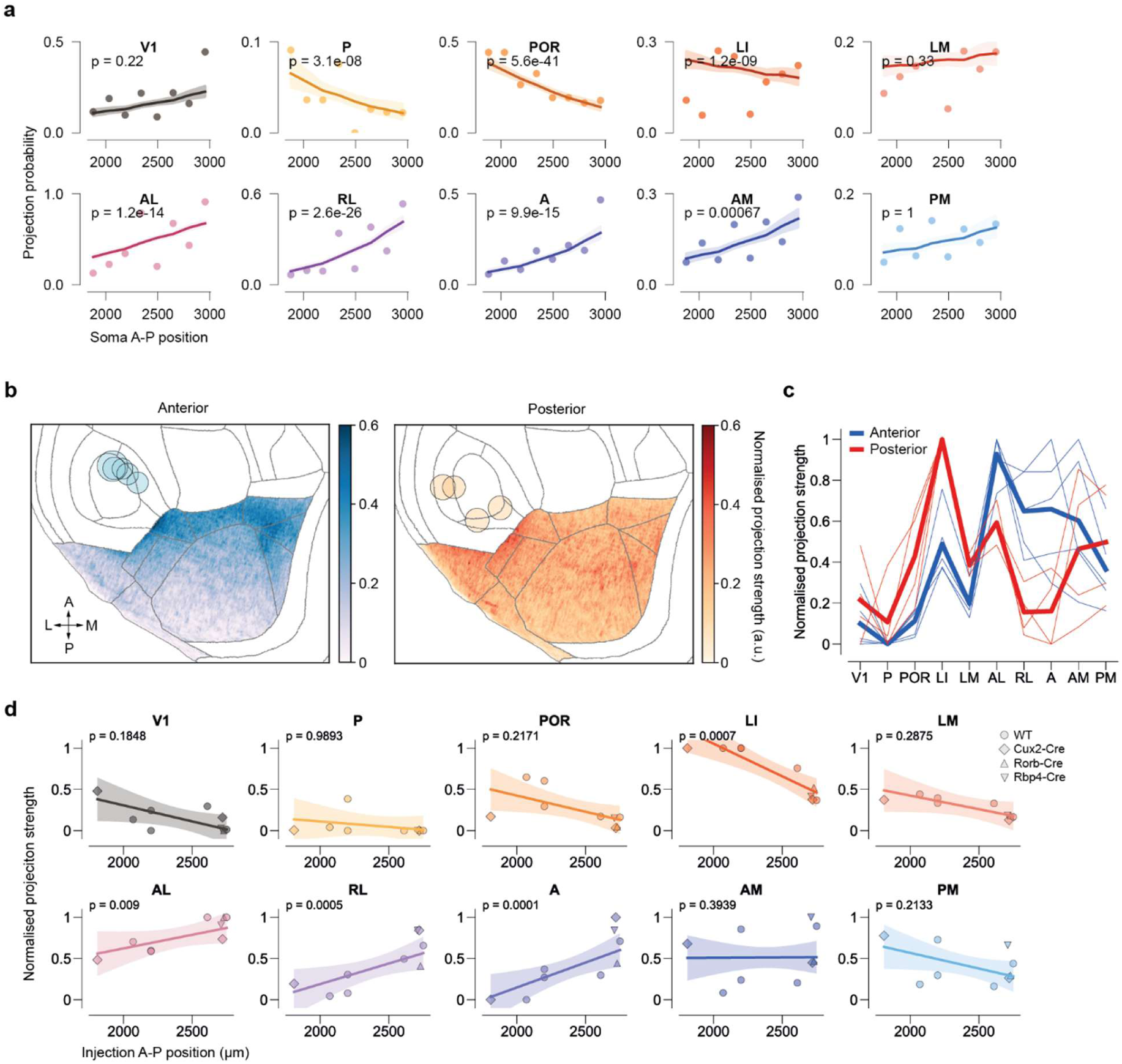
Anterior-posterior topographic mapping of A1 inputs to VC is not explained by mere proximity. **a,** Relationship between A-P source site position and targeting frequency (same as in Fig. 2g but displaying all visual areas). The y-axis shows the proportion of neurons targeting the given visual area for neurons in a particular A-P source site position. The x-axis is binned A1 source site A-P position with scale in microns normalised to most posterior VC position. P-values are calculated from logistic regression models including distance and mice as covariates. Fitted lines are predicted values from logistic regression model combining all mice together for A-P bin centres with mean distance for each centre as covariate, shaded area is 95% confidence limits.. **b,** Average normalised projection strength to visual cortex of neurons in anterior (n=5) or posterior (n=4) auditory cortex, computed from EGFP-injection data from the Allen Connectivity Atlas (Methods). Experiments were from WT (N=5), Cux2-Cre (N=2), Rorb-Cre (N=1) or Rbp4-Cre (N=1) mice. Data was normalised between 0 and 1 for each experiment before averaging. Circles indicate position of injection centroid, circle size corresponds to injection volume. **c,** Normalised projection strength to each visual area, for anterior and posterior injections. Thin lines correspond to individual experiments, thick lines indicate averages. **d,** Relationship between normalised projection strength and anterio-posterior position of the injection centroid for each visual area. Marker shape indicates transgenic line. Fit and p-values obtained with linear regression accounting for distance between injection location and visual areas, and inter-animal variability. P-values are corrected for multiple comparisons with the Bonferroni method.

**Supplementary Figure 5.**
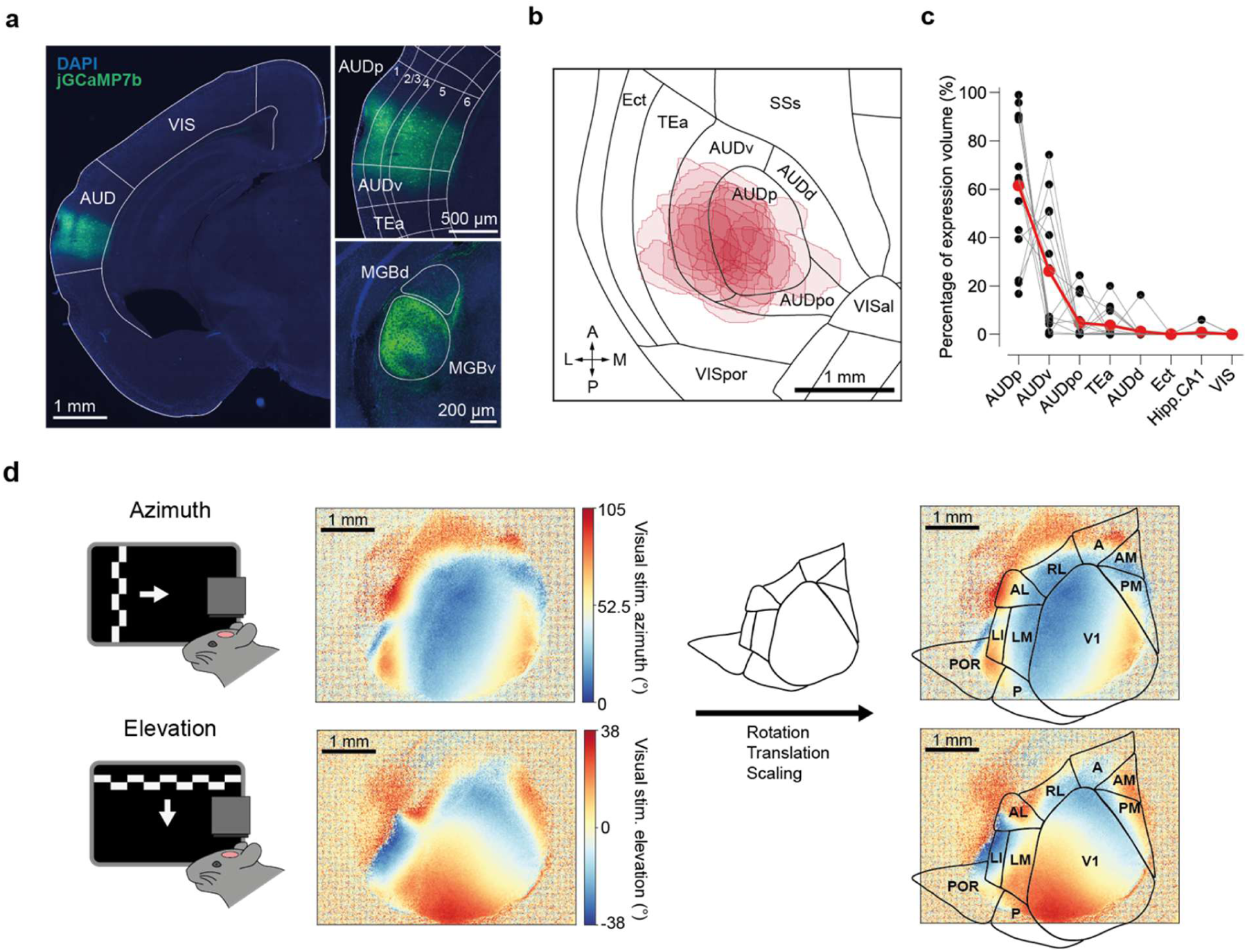
Auditory cortex injections for two-photon imaging and intrinsic imaging. **a,** Coronal brain slice at ∼bregma −2.9 mm (left, top right) or bregma −3.15 mm (bottom right), showing jGCaMP7b expression (green) in auditory cortex in an example mouse following stereotaxic injection. AUD: Auditory cortex, VIS: visual cortex. AUDp: primary auditory cortex; AUDv: ventral auditory cortex; AUDd: dorsal auditory cortex; AUDpo: posterior auditory cortex; TEa: Temporal association area. MGBv/d: Medial geniculate body, ventral/dorsal portion. **b,** Top view of jGCaMP7b expression area for each mouse (red areas), aligned to the Allen Brain Atlas. N= 15 animals. Ect: Ectorhinal cortex; SSs: Somatosensory supplementary area. **c,** Percentage of jGCaMP7b expression volume by brain area, in each animal (black) and averaged across animals (red). N = 16 animals. Hipp.CA1: Hippocampus, CA1 field. d, Retinotopic mapping using intrinsic imaging and area map alignment in an example mouse. Left: Schematic of visual stimulation with drifting gratings. Middle: Example azimuth (top) and elevation (bottom) retinotopic maps obtained with intrinsic signal imaging. Right: Alignment of visual area boundaries from Allen Brain Atlas to retinotopic maps, based on retinotopic reversal locations.

**Supplementary Figure 6.**
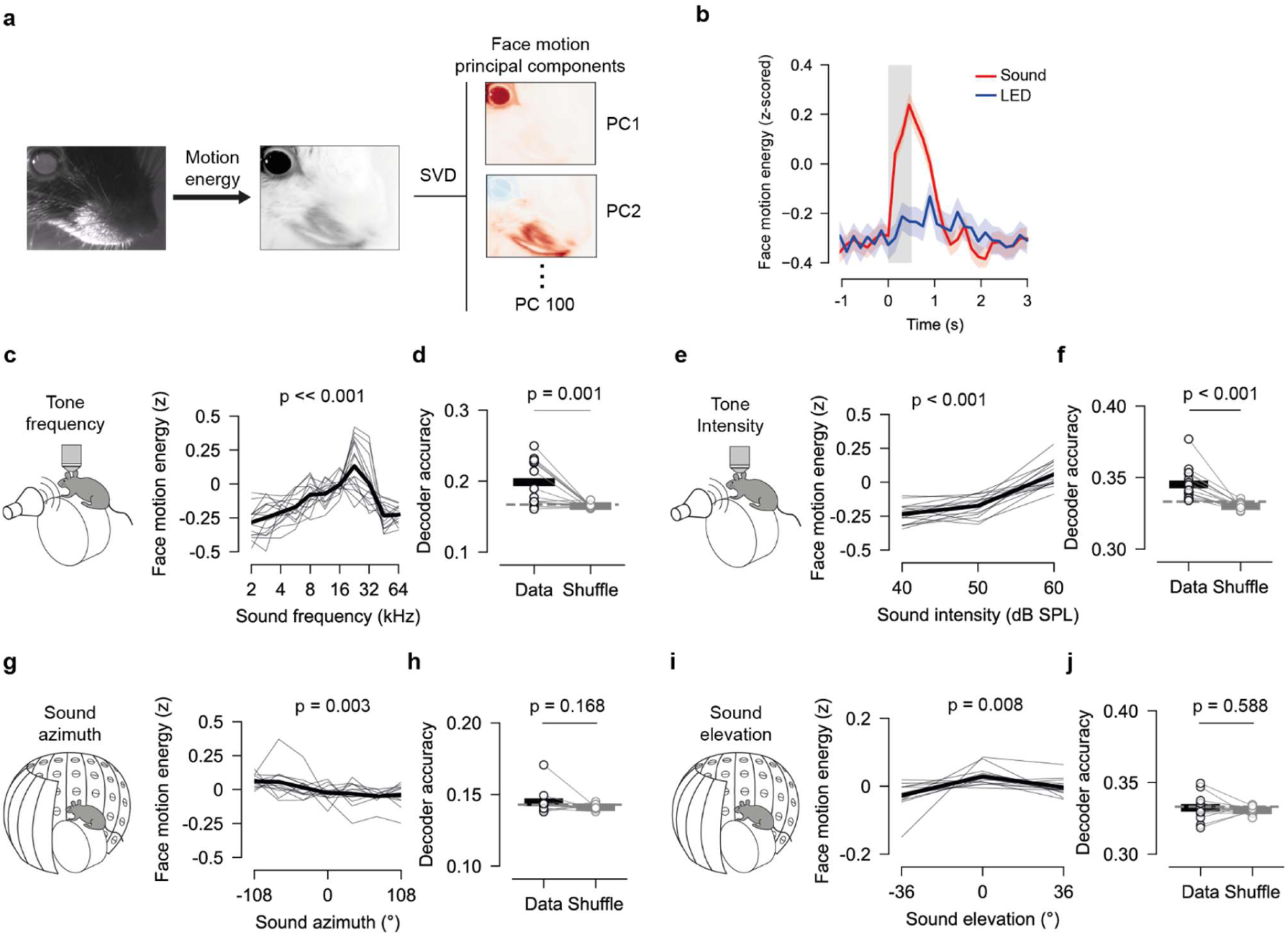
Mouse facial movements depend on auditory stimulus characteristics. **a,** Example frame from recording of mouse face during sound presentation. Motion energy was extracted from facial motion videos, and singular value decomposition (SVD) was used to obtain principal components of facial motion (Methods). **b,** Average face motion energy in an example session, aligned to the onset of auditory (red) or visual (blue) stimuli. Lines indicate mean ± sem. Shaded gray area indicates time of stimulus presentation. **c,** Relationship between tone frequency (left) and facial motion energy after stimulus presentation. Thin lines indicate average motion for one animal, thick black line indicates median across animals. P-value indicates significance of modulation with stimulus frequency (Friedman test). **d,** Accuracy of a support vector machine decoder (SVM) in estimating the octave of a pure tone based on the 100 first PCs of facial motion, using either measured or trial-shuffled data. P-value, Wilcoxon signed-rank test. **e-f,** Same as **c-d**, for pure tone intensity. **g-h,** Same as **c-d**, for sound azimuth. **i-j,** Same as **c-d**, for sound elevation

**Supplementary Figure 7:**
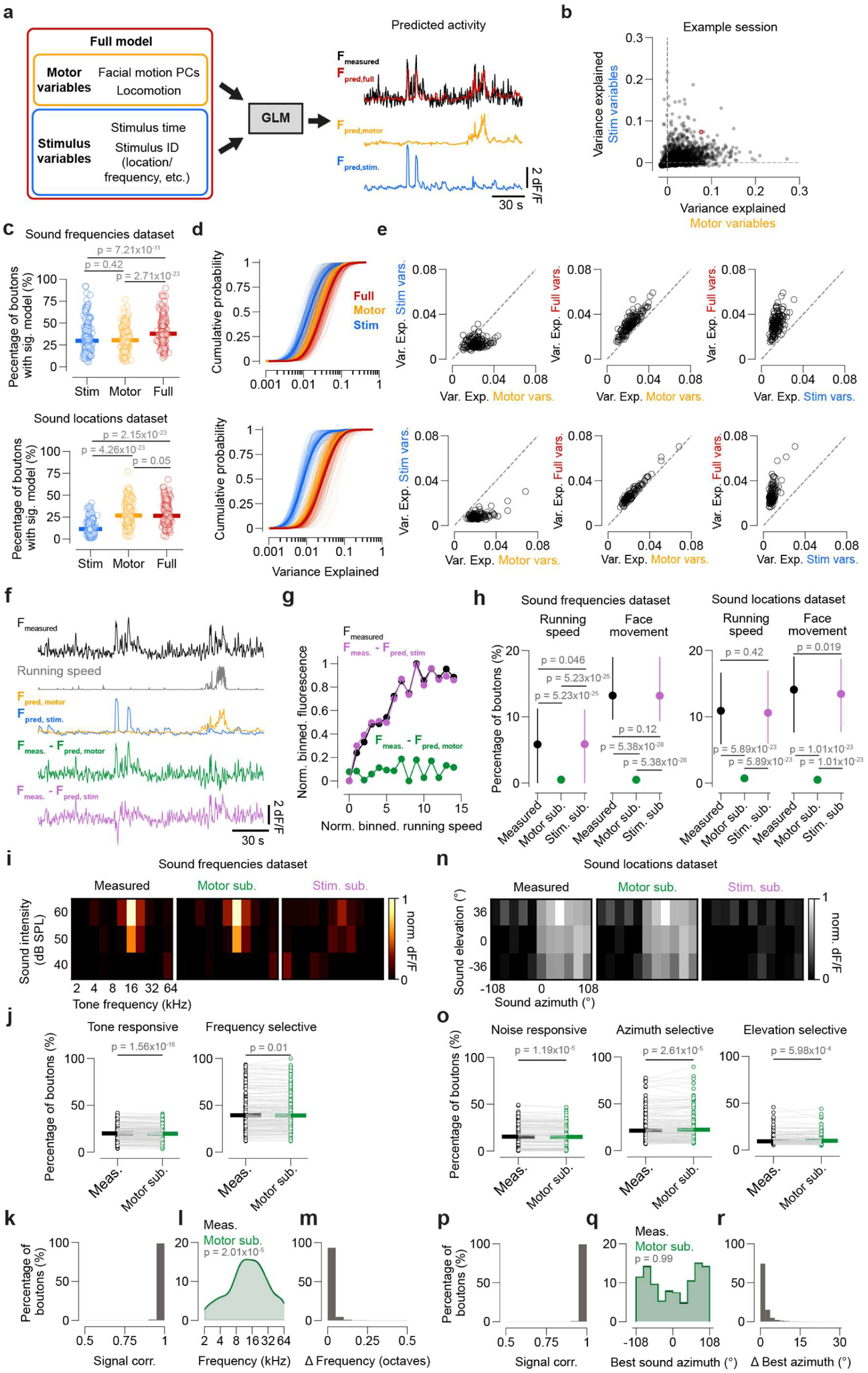
Subtraction of motor component estimated with generalised linear model minimally affects auditory representations in AC boutons. **a,** Left, schematic of GLM structure. A model using motor-related and stimulus-related variables was fit to the neural activity of each bouton. Right, example of an AC bouton activity trace (F, black), reconstructed using the fits from the stimulus-related, motor-related of all variables (F_pred,stim_, F_pred,motor_ and F_pred,full_, respectively). **b,** Cross-validated variance explained by motor variables vs stimulus variables for the AC boutons in one example session. Each point corresponds to one bouton. The traces in **a** correspond to the bouton with a red circle. **c,** Percentage of boutons in each session for which a model containing motor-related, stimulus-related or all variables explained a significant amount of variance across folds (Wilcoxon signed rank test), within the frequencies (top) or locations (bottom) dataset. Circles indicate proportion in one session, horizontal bars are median across recordings. N_frequencies dataset_=161 sessions, N_locations dataset_=135 sessions**. d,** Cumulative variance explained by stimulus-related, motor-related or all variables for all boutons with a significant model of the corresponding type, in the two datasets. Thin lines indicate the cumulative variance for boutons in one session, thick lines contain data from all sessions. **e,** Median variance explained by session for each set of variable combinations, in the two datasets. **f,** Example AC bouton fluorescence trace (F_measured_), normalised running speed (a.u.), fluorescence trace predicted by stimulus and motor models (F_pred,stim_ and F_pred,motor_) and trace after subtracting either the motor model prediction (F_meas. -_ F_pred,motor_) or the stimulus model prediction (F_meas. -_ F_pred,stim_). **g,** Example locomotion tuning curve of one AC bouton, computed from measured traces or from traces after subtracting predictions from motor variables (green) or stim variables (purple). Running was grouped into 15 equally spaced bins, points indicate the normalised avg. fluorescence at time points when locomotion speed was in a given bin. Error-bars indicate s.e.m. **h,** Percentage of boutons tuned to running speed or face movement (average percentage across face motion PCs) in each dataset, computed either from measured traces or from traces after subtraction of predictions from motor or stimulus variables. Points indicate median across recordings, error bars indicate 25^th^ and 75^th^ percentiles. **i,** Frequency-response areas (FRAs) of example AC bouton, computed from measured trace or from trace after subtracting motor or stimulus predictions. **j,** Percentage of tone responsive and frequency-selective AC boutons, computed from measured or motor-subtracted traces. Circles indicate percentage in one session, thick bars indicate median across sessions. **k,** Signal correlation between FRAs computed from measured or motor-subtracted traces. **l,** Distribution of best frequencies, calculated from measured or motor-subtracted traces. P-value from Wilcoxon signed rank test. **m,** Absolute difference (Δ) in best frequency calculated from measured or motor-subtracted traces. **n,** Location-response areas (LRAs) of example AC bouton, computed from measured trace or from trace after subtracting motor or stimulus predictions. **o,** Percentage of bandpass-noise responsive, sound-azimuth-selective or sound-elevation-selective AC boutons, computed from measured or motor-subtracted traces. **p,** Signal correlation between FRAs computed from measured or motor-subtracted traces. **q,** Distribution of best sound azimuth, calculated from measured or motor-subtracted traces. P value obtained with Kolmogorov-Smirnov test. **r,** Absolute difference (Δ) in best sounds azimuth calculated from measured or motor-subtracted traces. P values obtained with Wilcoxon signed-rank test expect where indicated otherwise

**Supplementary Figure 8.**
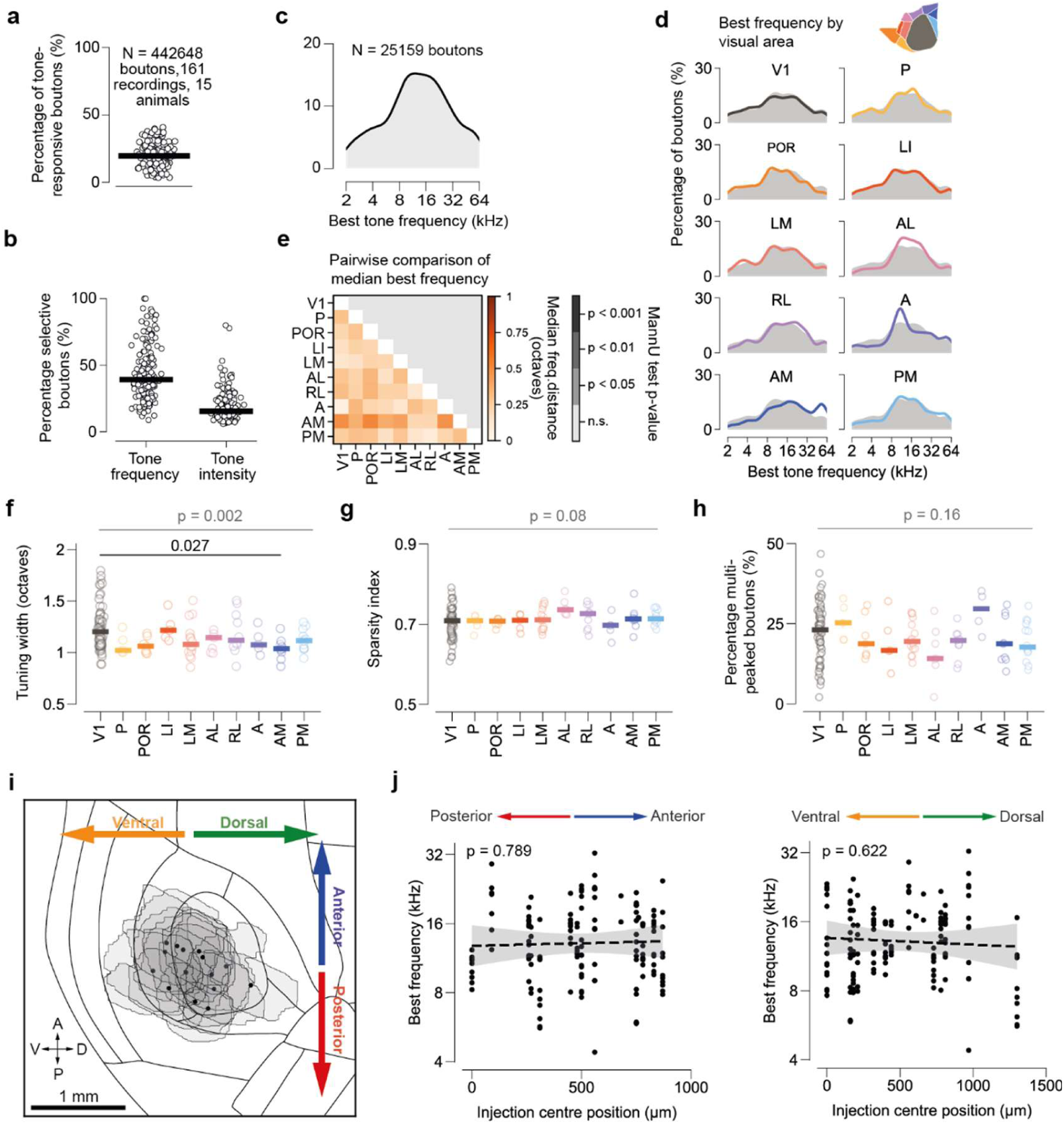
Comparison of tone frequency representations in AC→VC boutons across areas. **a,** Percentage of tone-responsive AC boutons in each recording. Black bar indicates median across recordings. **b,** Percentage of tone-responsive boutons that were selective for tone frequency or tone intensity in each recording. **c,** Distribution of best frequency for all single-peaked frequency-selective boutons. **d,** Distribution of best frequency for AC boutons innervating each visual area (filled distributions) or all areas combined (black line). **e,** Pair-wise comparison of median best frequency between visual areas. P-values obtained with hierarhical bootstrap. **f,** Median tuning width of frequency-selective boutons in each recording, grouped by area. Bar indicates median across recordings. gray P-value indicates comparison across all areas (mixed effects ANOVA), black p-value indicates comparison between area pairs (Mann Whitney U test). **g,** Median sparsity index (Methods) of frequency-selective boutons in each recording. **h,** Percentage of multi-peaked FRAs in each recording, grouped by visual area. See Fig. 3e for examples of single-peaked and multi-peaked FRAs. **i,** jGCaMP7b expression area for each mouse used for frequency-tuning experiments (gray areas), aligned to the Allen Brain Atlas. Black dots indicate injection centre positions. N = 15 animals. **j,** Relationship between anterio-posterior (left) or dorso-ventral (right) position of the injection centroid and best frequencies measured in axonal projections to visual cortex. Each point corresponds to one session. Fit and p-value obtained with linear regression

**Supplementary Figure 9.**
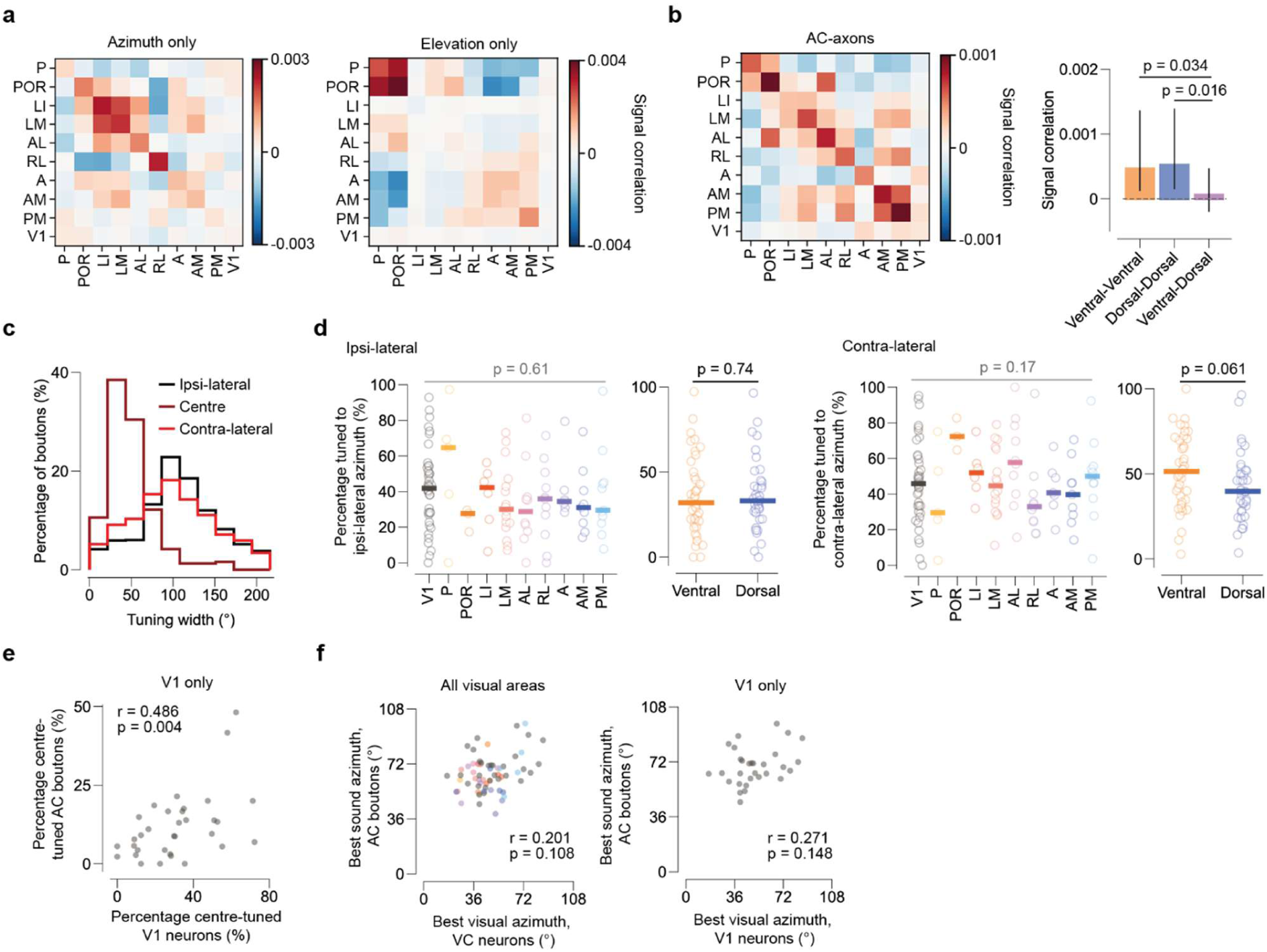
Additional analysis of sound location tuning in AC→VC boutons. **a,** Matrix of average pairwise signal correlations in azimuth (left) or elevation (right) tuning curves between AC boutons, grouped by visual area. **b,** Left, matrix of average pairwise signal correlations between location response areas (LRAs) of AC axons (i.e. after grouping boutons into axons based on activity correlations, see Methods). Right, signal correlation between axons of the same or different streams. **c,** Width of the azimuth tuning curve (full-width at half-max) for all azimuth selective boutons, grouped into ipsi-, centre-or contra-lateral tuned based on the peak of their tuning curve. **d,** Percentage of ipsi-lateral tuned (left) or contra-lateral tuned (right) boutons per recording, grouped by visual area and visual stream. P-values obtained with mixed-effects ANOVA. **e**, Correlation between percentage of centre-tuned V1 neurons and percentage of centre-tuned AC→V1 boutons in each spatial bin. R and p value obtained with Spearman correlation. **f,** Correlation between best visual location in VC neurons and best sound location in AC boutons in each spatial bin, for all VC (left) or within V1 (right).

**Supplementary Figure 10.**
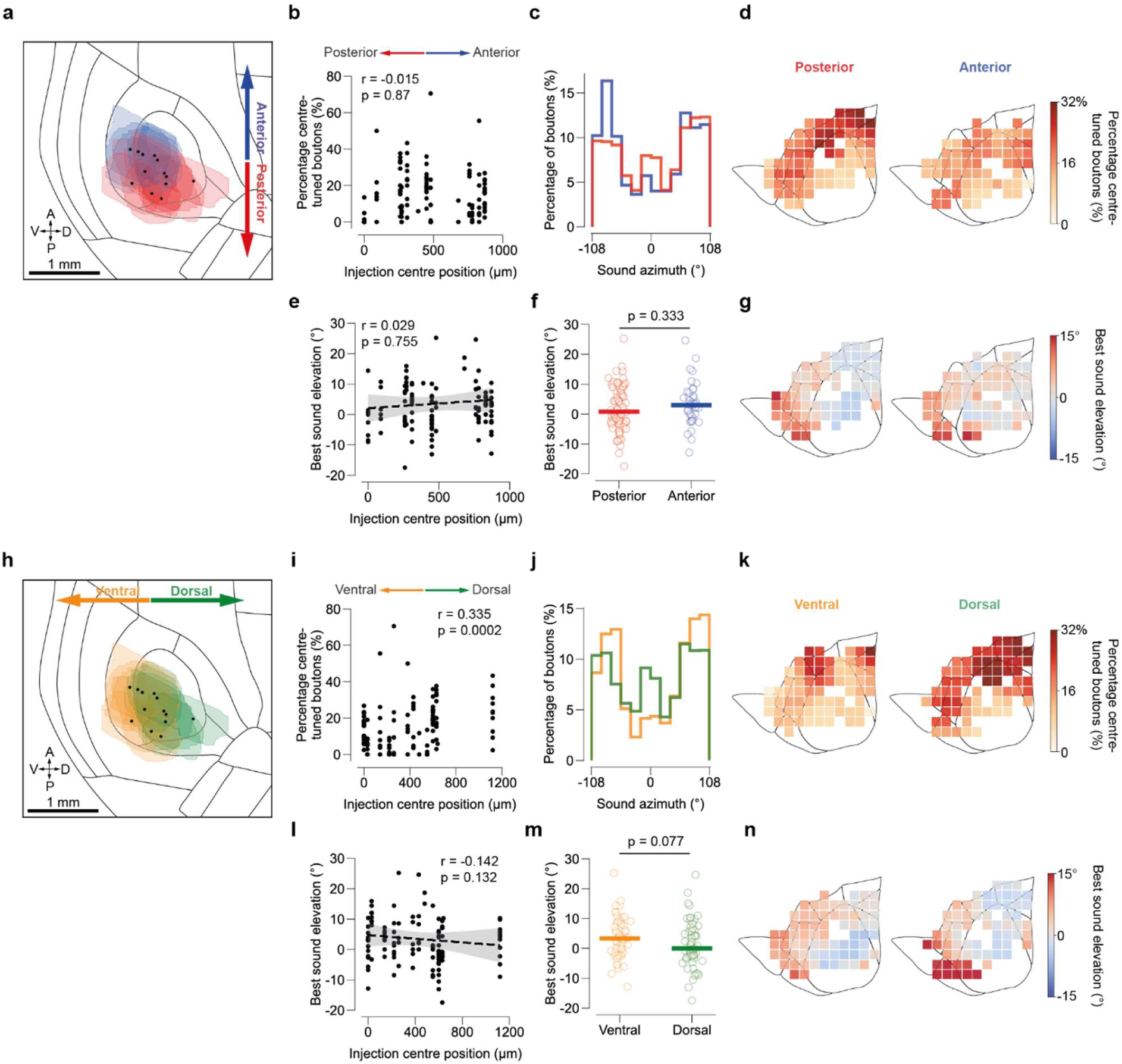
Effect of injection location on observed location tuning in AC→VC boutons. **a,** jGCaMP7b expression area for each mouse used for location-tuning experiments grouped into anterior (> 600 µm, N = 5) and posterior (< 600 µm, N =8) mice. Dots indicate injection centroids. **b,** Relationship between injection position in the antero-posterior axis and percentage of centre-tuned AC boutons. Each point corresponds to one session. R and p-value obtained with Spearman correlation. **c,** Distribution of best sound azimuth for boutons from animals with anterior (N= 2250 boutons) or posterior injections (N= 6601). **d,** Percentage of centre-tuned boutons per 300 µm bin in aligned cortical map, for animals with anterior or posterior injection location. **e,** Relationship between injection position in the antero-posterior axis and best sound elevation. Fit and p-value from linear regression. **f,** Average best elevation of AC boutons for each session, grouped by injection location. Lines indicate median across sessions. **g,** Best sound elevation per 300 µm bin on aligned cortical map for animals with anterior or posterior injection location. **h-n,** Same as **a-g**, comparing animals with ventral (< 400 µm, N =6) or dorsal (> 400 µm, N=7) injection sites. N_ventral, azimuth dist._: 3075 boutons; N_dorsal, azimuth dist._: 5776 boutons.

